# Dynamic design: manipulation of millisecond timescale motions on the energy landscape of Cyclophilin A

**DOI:** 10.1101/490987

**Authors:** Jordi Juárez-Jiménez, Arun A. Gupta, Gogulan Karunanithy, Antonia S. J. S. Mey, Charis Georgiou, Harris Ioannidis, Alessio De Simone, Paul N. Barlow, Alison N. Hulme, Malcolm D. Walkinshaw, Andrew J. Baldwin, Julien Michel

## Abstract

Proteins need to interconvert between many conformations in order to function, many of which are formed transiently, and sparsely populated. Particularly when the lifetimes of these states approach the millisecond timescale, identifying the relevant structures and the mechanism by which they inter-convert remains a tremendous challenge. Here we introduce a novel combination of accelerated MD (aMD) simulations and Markov State modelling (MSM) to explore these ‘excited’ conformational states. Applying this to the highly dynamic protein CypA, a protein involved in immune response and associated with HIV infection, we identify five principally populated conformational states and the atomistic mechanism by which they interconvert. A rational design strategy predicted that the mutant D66A should stabilise the minor conformations and substantially alter the dynamics whereas the similar mutant H70A should leave the landscape broadly unchanged. These predictions are confirmed using CPMG and R1ρ solution state NMR measurements. By accurately and reliably exploring functionally relevant, but sparsely populated conformations with milli-second lifetimes *in silico*, our aMD/MSM method has tremendous promise for the design of dynamic protein free energy landscapes for both protein engineering and drug discovery.

## Introduction

Capturing the complete dynamics of proteins to understand their many functions has long been a goal of structural biology. Experimentally, trapping proteins into relevant conformational states and characterising these at atomic resolution provides tremendous insight^1–3^. Such static ‘snapshots’ are inherently incomplete and especially in cases where proteins are particularly dynamic, understanding function can require characterisations of conformational states that are formed only transiently, and populated as sparsely as a few per cent^4–6^. These conformers are typically challenging to characterise experimentally. Solution-state NMR has proven unique in its ability to detect and structurally characterise these functionally relevant states, whose lifetimes can be on the order of a few milliseconds and are otherwise ‘invisible’ to experimental measures^6–8^. Such states can play important roles in processes as diverse as protein folding, molecular recognition and catalysis, but remain challenging to characterise ^9–16^. To engineer proteins with new functions, and for drug discovery, there is a clear need to be able to characterise, explore and manipulate the full set of conformational states accessed by proteins, and the mechanism by which they interconvert.

Molecular dynamics (MD) simulations provide an attractive means to accomplishing this goal *in silico*. To characterise low amplitude structural fluctuations in local minima, simulations on timescales of a few hundred nano-seconds may be sufficient^17,18^. Substantially longer simulations are required in order to sample sparsely populated ‘excited’ conformational states with millisecond lifetimes. Such a ‘brute force’ calculation is possible at present only for relatively small proteins using dedicated hardware that is not widely available^19^. To make progress, two broad categories of enhanced sampling methods have been developed with the goal of increasing total effective time covered by a simulation. Predefined collective variables can be exhaustively sampled (Metadynamics^20–22^, Umbrella Sampling^23–25^, Steered MD^26–28^), or the potential energy function of the forcefield can be explicitly altered (scaled MD^29–32^ accelerated MD (aMD)^33,34^). The former class of methods require the degrees of freedom underpinning the motion of interest to be supplied in advance, which pre-supposes that the relevant motions are already known. The latter class of methods can overestimate the population of high energy regions of the free energy surface (FES) and so explore unphysical regions of conformational space.

Aiming to improve upon these methods, Markov State Modelling (MSM) based sampling approaches have emerged as an exciting method for sampling protein dynamics on millisecond timescales^2,35–41^. These techniques may be used to sample conformational space using a series of relatively short MD trajectories. These trajectories are analysed with the goal of obtaining a discrete set of interconverting conformational ‘states’, where each has no knowledge of any previously visited state. This method allows a series of independently generated MD trajectories to be combined into a complete description of the energy landscape of the protein. MSM have been shown to reproduce experimental observables in model systems where the structures of the relevant states were known in advance^35,42,43^, and in cases where the timescales for conformational transitions were sufficiently short to be validated with ‘brute force’ MD methods^19,37,44^.

In this work, we generate a protocol suitable for sampling transitions between structurally ambiguous states separated by millisecond kinetic barriers (Fig. 1b). This is achieved by harnessing the ability of aMD to efficiently explore conformational space, while relying on the MSM procedures to recover a more accurate thermodynamic and kinetic information description of the system. Because this approach provides atomically detailed structural ensembles and detailed information about how they interconvert, it is well suited to both generate experimentally testable mechanistic hypotheses and enable the rational design of conformational ensembles.

**Figure 1.**
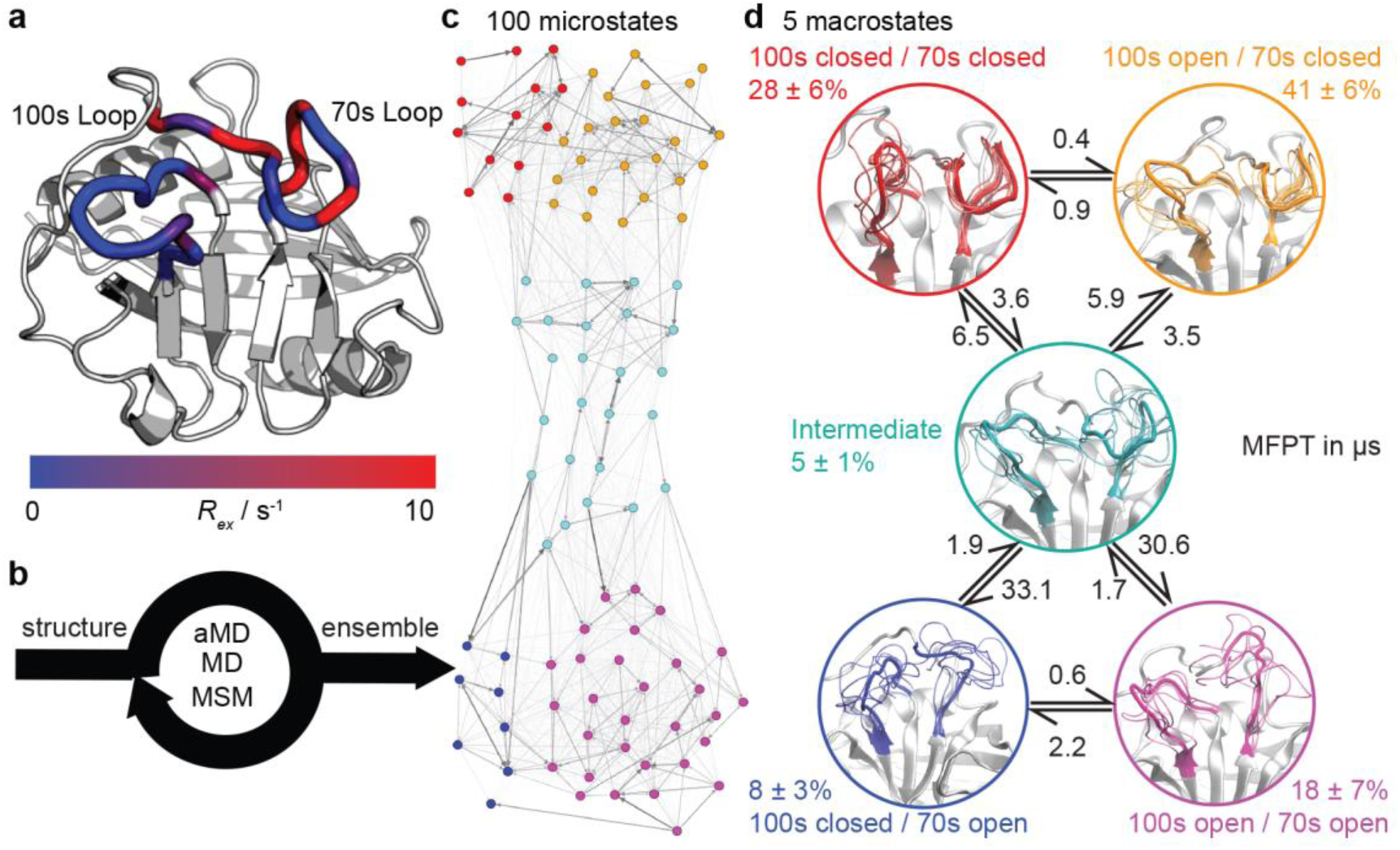
Calculation the dynamic ensemble of WT CypA. **a** an X-ray diffraction structure of CypA, with the 100s and 70s loops indicated. A number of residues have been previously determined to undergo conformational exchange^68,69^ which is evidence by a large value o f *R*_ex_ for these residues, measured by N MR. These values are consistent with the formation of transiently sampled, but sparsely populated alternative loop conformations. **b** It is desirable to determine the nature of the interconverting structures, their populations and mechanism by which they exchange. We accomplish this for CypA using a combination of aMD, MD and MSM methodologies (detailed methods). **c** A 100 microstates MSM for wild-type CypA that describes 70s and 100s loop motions. The width of the arrows is proportional to the inter-conversion rates. **d** The microstates were clustered into a mini mal set of sub-states that together contain the relevant amplitudes of motion and time-scales of state-to-state inter-conversion present in the full ensemble. The calculated rates and populations are indicated. Error bars on reported populations were obtained by bootstrapping of the MD trajectories assigned to the individual microstates.

We test the novel aMD/MSM protocol by both characterising and rationally modifying the free energy surface of human cyclophilin A (CypA). This enzyme is a human prolyl isomerase whose incorporation into new virus assemblies is essential for HIV-1 infectivity^45–48^ and HCV replication^49^, making it a major drug target^50–57^. This protein has been widely studied^58–67^ and possesses two functional loops, residues 65-77 (70s loop, Fig. 1a) and 100-110 (100s loop, Fig. 1a) that have been established by NMR to undergo substantial dynamics on the millisecond timescale^68–70^. An atomistic picture of the conformational changes has proven elusive. Our aMD/MSM procedure reveals a range of conformational sub-states of CypA where both the 100s and 70s loops exchange between open and closed states (Fig. 1c/d).

To manipulate the energy surface, we implemented a design strategy based on hydrogen bonding patterns to stabilise specifically sparsely populated conformers with millisecond lifetimes (Fig. 2). This procedure led us to a previously unidentified mutation D66A. Whereas WT has the 70s loop predominantly in the closed state, consistent with the majority of X-ray structures of CypA solved to date^52,55,70–75^, D66A was predicted to predominantly occupy the open state of the 70s loop. Similarly, as a negative control, a second similar mutation H70A was predicted by aMD/MSM to closely resemble the WT (Fig. 3).

**Figure 2.**
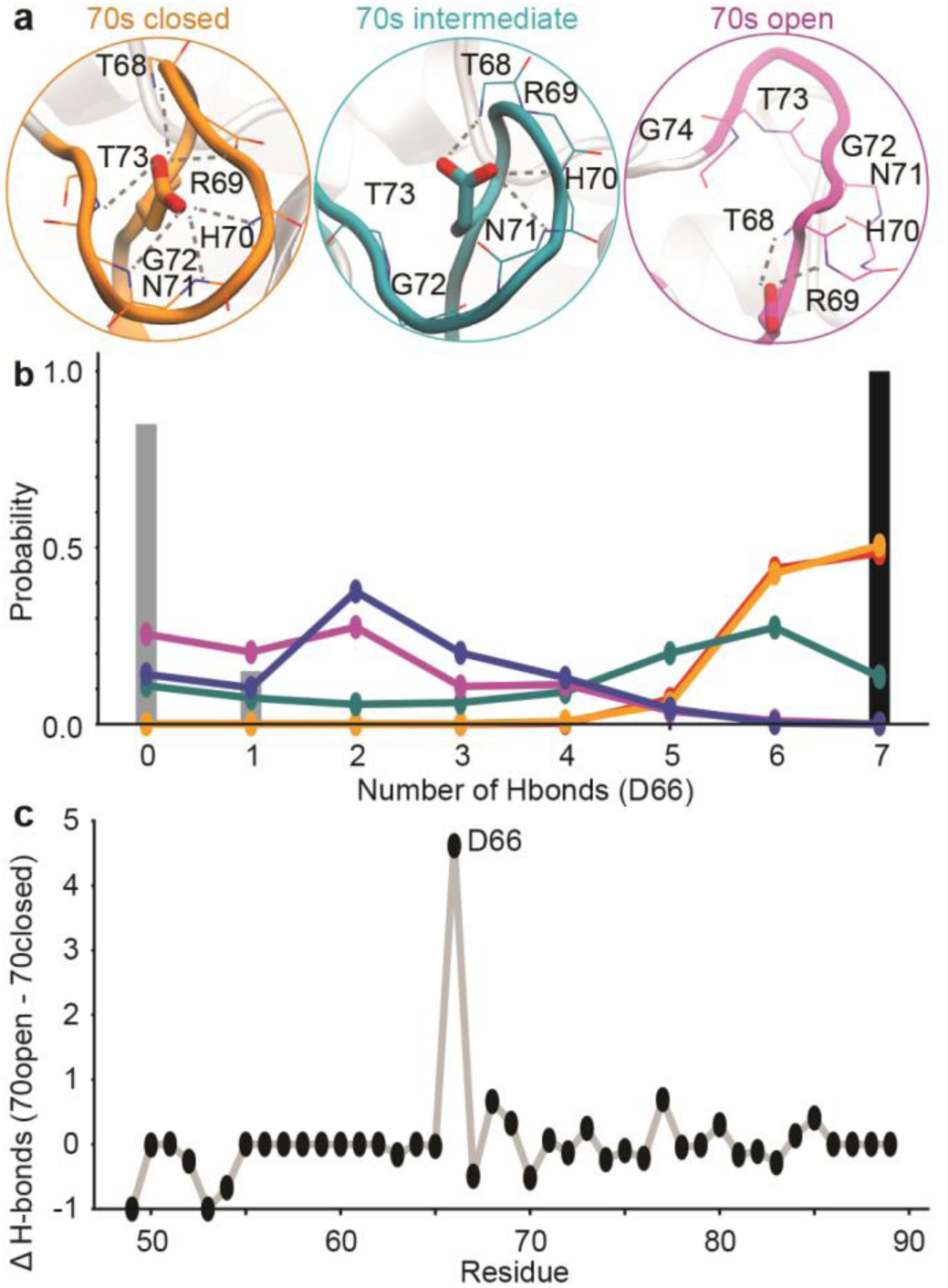
Design of ensemble disrupting mutation. **a** Specific interactions made by D 66 with 70s loop residues in representative MD snapshots of the closed, intermediate and open states. **b** The probability distribution of the numbers of intra-molecular H-bonds between D66 and 70s loop residues in the 5 sub-states. These are compared to X-ray structure PDB 1AK4^71^ (black), and NMR ensemble PDB 2N0 T (grey)^61^. **c** The difference in the average number of intra-molecular H-bonds in the 70s closed (orange/red) and 70s open (blue/purple) states. Residue D66 stands out as ‘ designable’ as it has substantially more H -bonds stabilising the closed state than the open state of the 70s loop.

**Figure 3.**
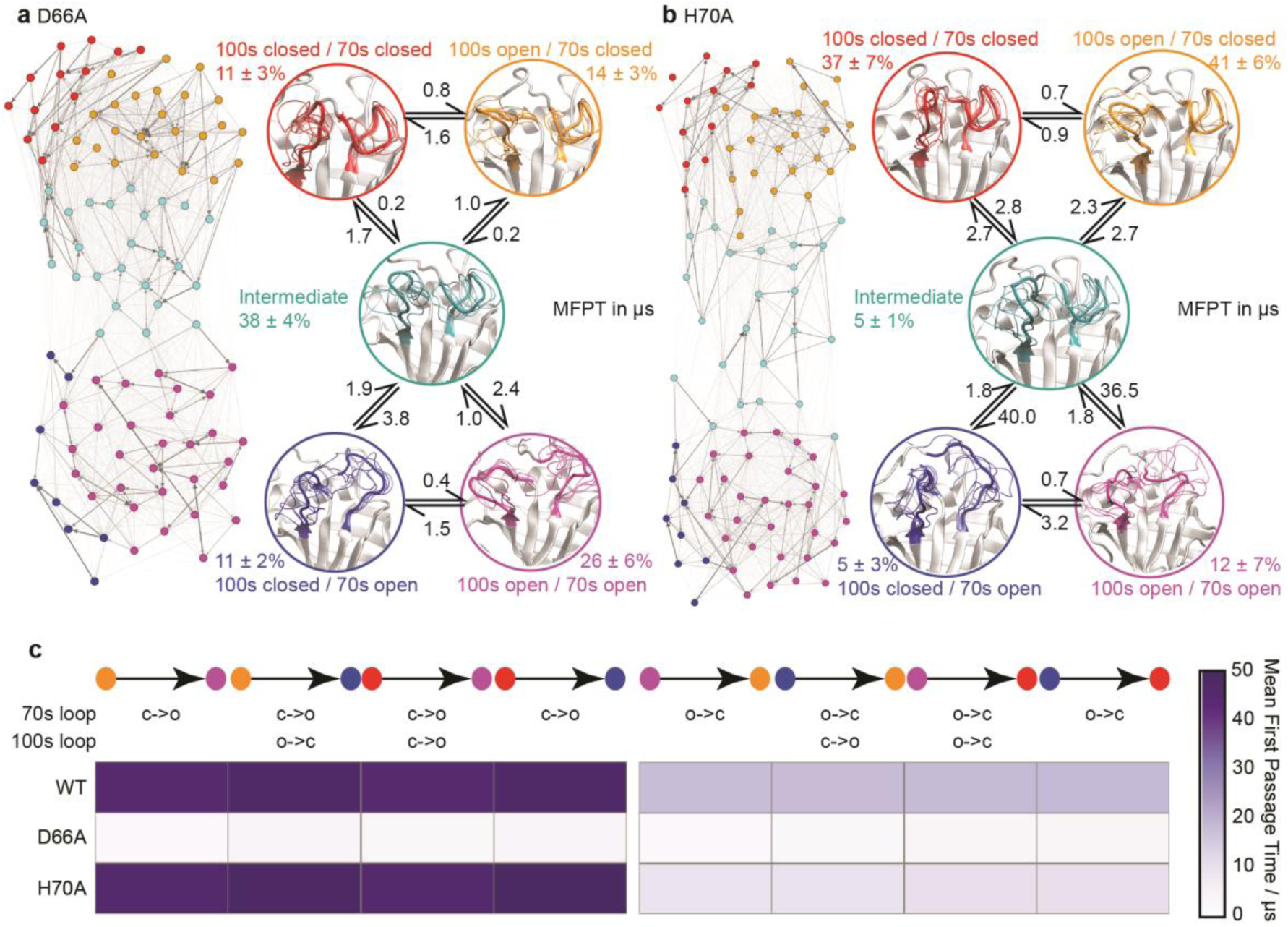
Ensembles of mutant CypA proteins D66A and H70A. **a** MSM model for D66A. **b** MSM model for H70A. Other details as in Fig 1c. **c** Heatmap of calculated mean first passage times (MFPT) between non-adjacent macro-states. The symbols ‘ c’ and ‘ o’ denote transitions between closed and open loop conformations.

Using a combination of R_1ρ_ and CPMG NMR experiments, with X-ray crystallography we tested these predictions (Fig. 4,5) and verified that exactly as predicted, the 70s loop is substantially disordered and open in D66A, and adopts a conformation very similar to the minor conformations populated in WT ensemble. Also consistent with the predictions of aMD/MSM, the free energy landscape of H70A was largely the same as for WT.

**Figure 4.**
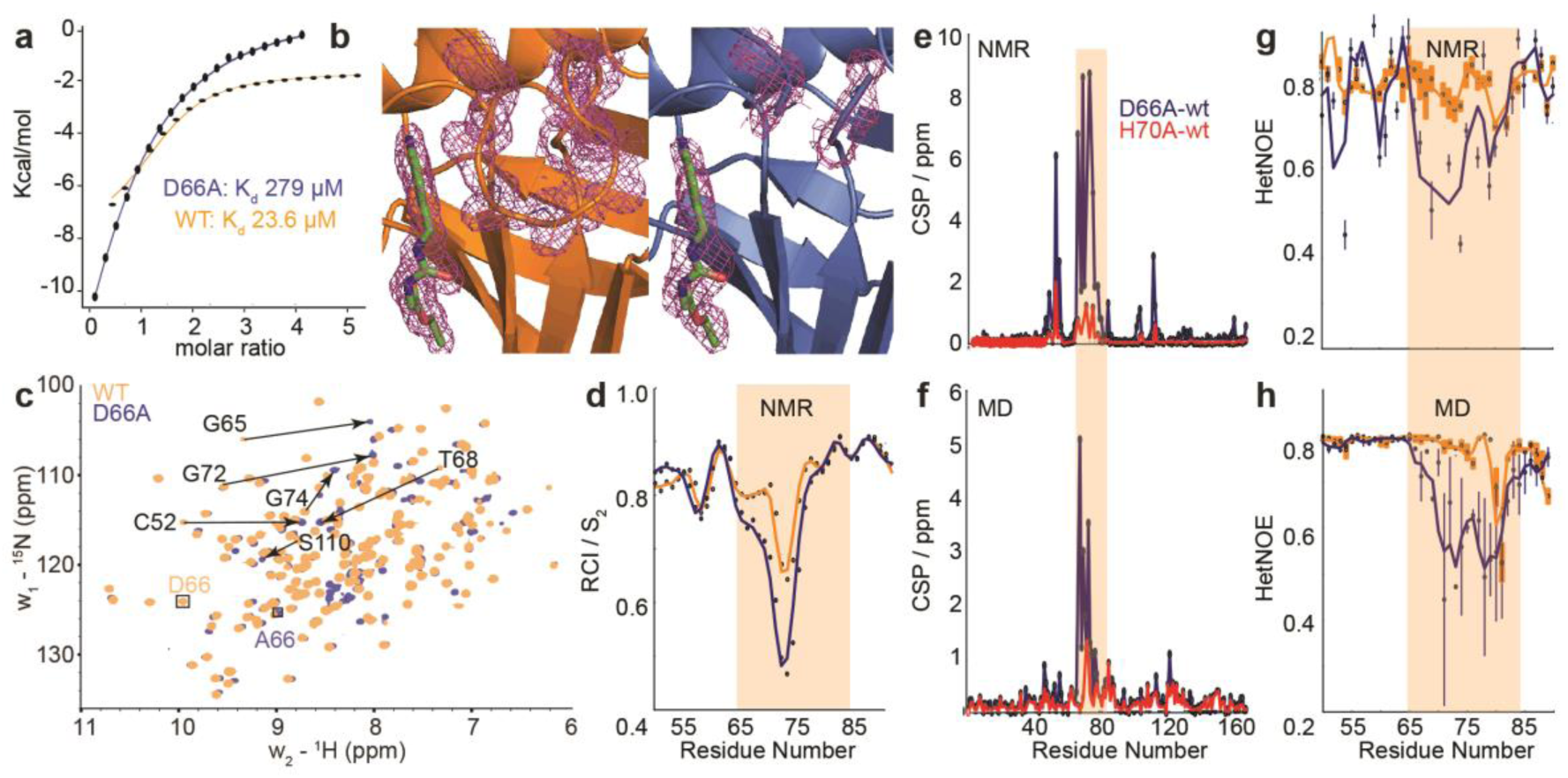
Comparison of biophysical characterisation and MD/ MSM conformational ensembles for WT CypA, D66A and H70A. **a** Binding isotherms for W T and D66A measured by ITC against an inhibitor. **b** X-ray structure of WT and D66A bound to inhibitor. Magenta meshes indicate 2 *F*o-*F*c electron density map at an isocontour of 1.0*σ* for the inhibitor and protein residues 65 to 75. **c** ^1^H-^15^N HSQC correlation spectra of WT and D66A at (10 °C), residues showing significant CSPs are highlighted. **d** Random coil index S_2_ values calculated from Talos+ for W T and D66A revealing increased disorder in the vicinity of the 70s loop for D66A. **e** Measured CSP per residue with respect to WT for D66A (blue) and H70A (red)^69^. **f** Calculated CSPs per residue from the computed ensembles. **g** Measured steady-state ^15^N-{^1^H} heteronuclear NOE transfers for W T and D66A in the vicinity of the 70s loop. **h** Calculated ^15^N-{^1^H} heteronuclear NOE transfers in the vicinity of the 70s loop from the computed ensembles.

**Figure 5.**
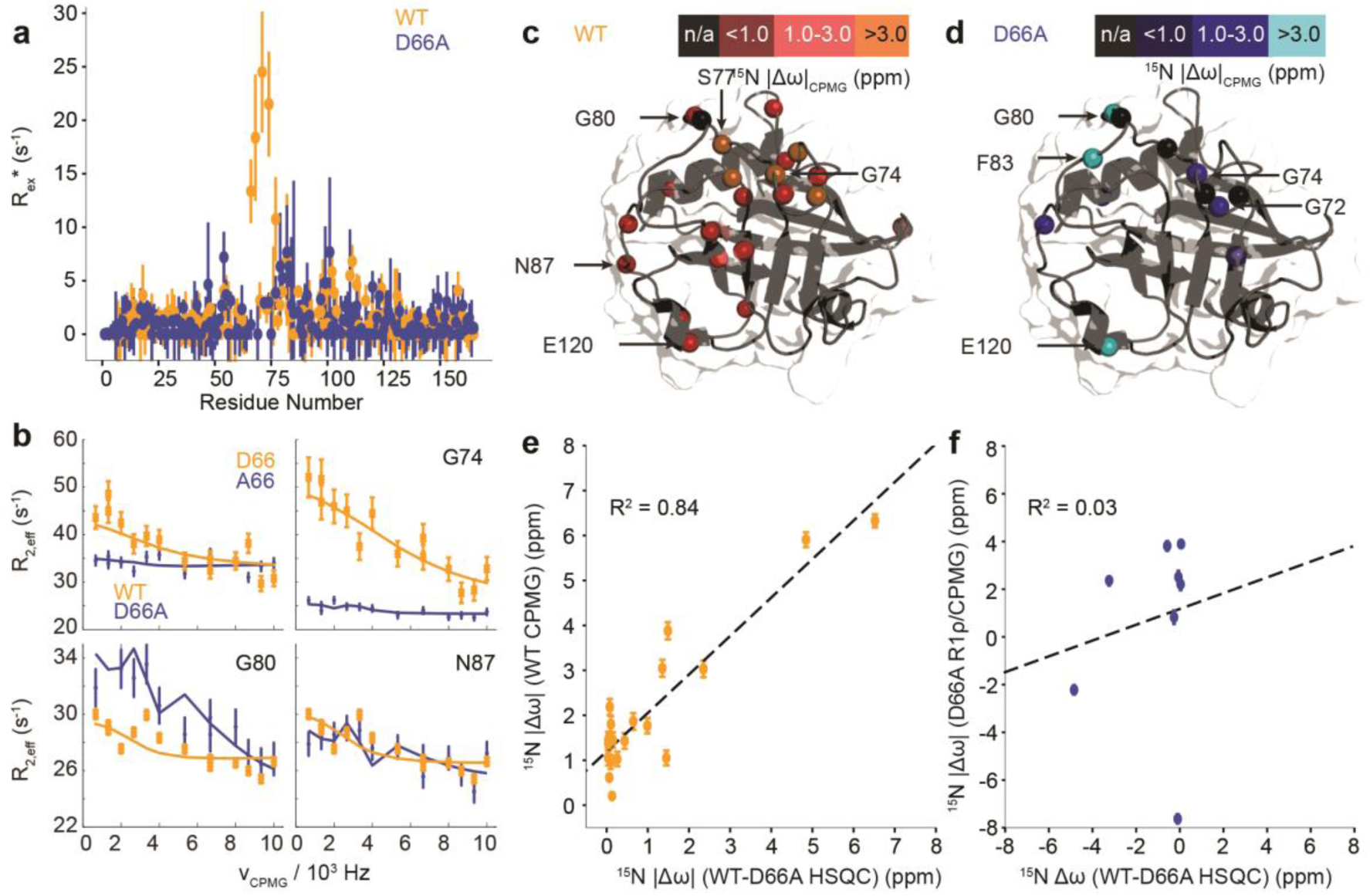
Characterisation of μs-ms dynamics in CypA and D66A by CPMG and R1ρ NMR measurements. **a** estimated contribution to relaxation from exchange, taken as the average of the difference between the lowest and highest frequency measurements from CPMG dispersion curves. **b** Comparison of selected dispersion profiles measured for CypA (orange) and D66A (blue) at 283 K and 600 MHz. **c** The experimentally determined chemical shift changes, Δ*ω*, from analysis of the CPMG/R1ρ curves for W T and (**d**) D66A. **e** Comparison of magnitude of Δω between CypA and D66A from HSQC measurements (x-axis) and from CPMG/R1ρ measurements on the WT (y-axis). The agreement between the two is remarkably good. **f** Comparison of magnitude of Δ*ω* between CypA and D66A from HSQC measurements (x-axis) and from CPMG/R1ρ measurements on D66A (y-axis). The agreement is poor revealing that the ‘ excited state’ in D66A is not the ground state of WT CypA.

Overall, we demonstrate that the aMD/MSM method accurately predicts the conformational fluctuations of the challenging and dynamic protein CypA including states with millisecond lifetimes. The resulting free energy surface is sufficiently accurate to enable the design of mutants that stabilise specific conformations. aMD/MSM provides a powerful tool for not only the atomistic characterisation, but also the manipulation of the free energy surface of dynamic proteins, including controlling states with millisecond lifetimes.

## Results

### Atomic-level insights in millisecond timescale CypA loop motion mechanisms

All-atom accelerated molecular dynamics (aMD) simulations ^33,34^ were initiated from an X-ray structure of wild type CypA (PDB ID 1AK4^71^). In aMD conformational sampling is accelerated by adding a biasing potential to the molecular mechanics potential energy function when it falls below a threshold (supplementary methods). Structures sampled from the aMD trajectory were used to seed sets of equilibrium MD trajectories, which were used for Markov state model (MSM) analysis. In a MSM, the protein dynamics is modelled as a set of Markov jump processes between discrete conformational states. These Markov processes can be thought of as jumps between nodes in a network, with the edges representing transition probabilities between these nodes, and where no knowledge of previously visited nodes is required to estimate the probability for a particular transition. The procedure enables reconstruction of a conformational ensemble from sets of independently run short MD trajectories that have each individually explored only a small portion of that conformational ensemble. Three rounds of MD simulations and MSM estimations were required to produce a 100 microstate MSM (Fig 1c) that was numerically stable, corresponding to a cumulative MD sampling time of 187.5 μs. Details about model validation against experimental observables are given in the Supporting Information (Figure S1-S3).

To obtain physical insight into the calculated ensemble, the 100 MSM microstates were further lumped into a coarser 5-macrostate model (see SI methods). This five state conformational landscape reveals that both the 100s and 70s loops can adopt ‘open’ and ‘closed’ conformations. While the inter-conversion of both loops were largely independent, the inter-conversion of the 100s loop was one order of magnitude faster than the 70s loop (Fig 1d, Fig S4), suggesting that the slower loop dynamics of the protein are largely associated with the opening and closing of the 70s loop. The most populated state for WT CypA, (100s open/70s closed, Fig. 1c/d orange, 41±6%) closely resembles the conformation typically observed in X-ray structures of CypA (Fig. S4), and fluctuations within the state were small (Cα RMSF values of 2.7 Å and 1.1 Å for the 100s and 70s loops, respectively). From this state, the 100s loop can rapidly open to adopt the 100s closed/70s closed state (Fig 1c/d, red, RMSF 2.6 Å and 1.1 Å).

These two conformational states must overcome a substantially larger kinetic barrier to adopt open conformations of the 70s loop (100s open/70s open, magenta) and (100s closed/70s open, blue). In contrast to the 70s closed states (red/orange), the 70s loop in its open conformations is relatively disordered, as evidenced by RMSF values >4 Å. These two 70s open states can interconvert, with the 100s loop opening/closing in an analogous fashion to the inter-conversions observed in the 70s closed states. While fluctuations of the 100s loop are rapid and largely independent of the 70s loop conformations, the opening/closing transition of the 70s loop was more complex, passing via an intermediate fifth state (teal) where both 70s and 100s loops display positional fluctuations that are intermediate between open and closed (RMSF ca. 2.9 Å). Qualitatively, it is important for the 100s loop to simply ‘not be in the way’, as some conformations of the open 100s state can effectively block the opening of the 70s loop.

To validate these findings we sought a mutation that would stabilise the group of minor, 70s open, conformations (teal/blue/purple). The number of hydrogen bonds each residue makes to the 70s loop was assessed in both the open and closed forms (Fig. 2b). Notably, D66 was found to adopt a large number of H-bonds in the open form, and substantially fewer in the closed (Fig. 2c), making this residue stand out as ‘designable’ and provide a target for altering the free energy landscape of CypA. The critical role of D66 in the proposed 70s loop motions mechanism is not supported by a recent NMR structural ensemble that has been proposed to describe the CypA 70s loop conformational changes measured by CPMG experiments.^61^ Analysis of the structures in that NMR bundle shows little changes in hydrogen bonding interactions with the 70s - loop (Fig. 2b). Further none of the structures in the NMR bundle contain a 70s -loop conformation similar to that commonly observed in high-resolution X-ray structures of CypA (Fig. S5). aMD/MSM simulations of D66A were performed as described for WT CypA (Fig 1). The resulting ensemble predicted that the population of the previous ground state was substantially reduced (from ca. 40% to ca. 15%, Fig. 3a), and the intermediate (teal) state was increased in a compensatory fashion to become the new ground state (from ca. 5% to ca. 40%). Overall with respect to CypA the population of states in which the 70s-loop is closed/open transitioned from ca. 70:30% to 25:75%. Moreover, the mean first passage times between the conformational states were predicted to decrease by 10-15 fold with respect to wild type (Fig. 3a and 3c). Finally, as a negative control, we performed aMD/MSM simulations on the mutation H70A^69^. The hydrogen bonding pattern for this residue was predicted to be broadly similar when the 70s loop is either open or closed (Fig. 2c). Consistent with this, aMD/MSM simulations determined that the conformational ensemble for H70A should resemble WT (Fig. 3b).

### Experimental validation of the calculated ensembles

To establish the validity of mechanistic hypotheses originating from the aMD/MSM models, we sought to compare our results to experimental data. Initially, we examined binding of an inhibitor against both WT and D66A. The *K*_d_ for D66A was 12-fold weaker than for WT (Fig. 4a), indicating a substantial change in the underlying protein conformations. Crystallisation of D66A proved challenging as we might expect for a protein with increased disorder, although both WT and D66A were successfully crystallised in the presence of the inhibitor^51^ and their structures were solved (Table S2) Comparison of electron density profiles revealed that while the 70s-loop adopts as expected a closed conformation in the WT complex (Fig 4b orange), electron density in the 70s loop for the D66A complex was largely absent (Fig. 4b blue), consistent with the notion that this loop occupies a predominantly disordered conformation in D66A.

To probe the conformational dynamics more directly, isotopically enriched samples of WT and D66A CypA were prepared for NMR analysis (Fig. S6-S7). ^1^H-^15^N HSQC spectra of both proteins revealed that while both were predominantly well structured (Fig. 4c, Fig. S8), there are substantial differences in the ground-state ensembles, exactly as predicted from the simulations. On a per-residue basis, the largest deviations in observed chemical shift positions between D66A and WT were found in the 70sloop region (Fig. 4e). The changes in chemical shift were quantitatively estimated by calculating the chemical shifts of individual conformers in the aMD/MSM ensembles using the program shiftx2^76^. The calculated CSP values are consistent with the experimental observation of significant changes in the 70s loop region, though the magnitude of the CSPs is underestimated (Fig 4f).

To determine if there are any changes in fast timescale motions of the 70s loop, the random coil index (RCI) was calculated for WT and D66A from the observed chemical shifts (Fig. 4d). These confirmed that the 70s loop is substantially more disordered in D66A, as expected from the simulations. This finding was further corroborated by measurement of the H-N steady-state heteronuclear NOE, where substantially lower values were recorded in the 70s loop for the D66A versus the WT. The heteronuclear NOEs were calculated from the MD ensembles, and also found to be in reasonable qualitative agreement with the experimental measurements (Fig. 4g/h).

To further characterise the fast timescale dynamics, *R*_2_ and *R*_1_ values were additionally measured at two magnetic field strengths, and a model-free analysis was performed. Such an analysis requires successively more complex motional models to be fitted to individual residues, in order to achieve an overall self-consistent picture of the dynamics. The procedure was unable to provide a self-consistent picture of relaxation for either CypA or D66A, with large variations in parameters between adjacent residues and ‘fast’ local fluctuations measured to be comparable to the global tumbling rate (Fig. S9). This is likely because of the substantial contributions of chemical exchange to the individual measurements (*vide infra*). Of the measurements obtained, the heteronuclear NOE nevertheless strikes a reasonable compromise between being relatively resistant to the effects of chemical exchange, whilst still being sensitive to relatively rapid local fluctuations within the molecule (Fig. 4g/h).

While the fast ps/ns time-scale dynamics indicate substantial differences in the 70s loop, the major goal was to determine if the structure of D66A resembles the excited state of WT CypA, thus validating the design strategy. A combination of R_1ρ_ and CPMG experiments were employed to study these conformational fluctuations on the milli-second timescale (Fig. 5a,b). The CPMG data for WT strongly resembles that obtained previously^68,69^. The combined data were analysed globally within a modified version of the software CATIA^77^. Such an analysis can ascertain a minimally complex binding model, and provide experimental measures of the kinetic and thermodynamic parameters that govern the exchange. Moreover, the analysis also provides chemical shifts (Fig. 5c,d), which characterise the structural changes that accompany the millisecond conformational rearrangements. For WT, a two-state model was found to explain well the combined dataset to within experimental error. The exchange rate *k* _*ex*_ was determined to be approximately 2200 s^-1^ and a population of excited state was determined to be approximately 2 % (Table S3).

Remarkably, the chemical shift differences directly measured from WT and D66A HSQC spectra were in good agreement with the chemical shift differences between ground and excited states of WT obtained by CPMG/R1ρ measurements (Fig. 5e). This correlation strongly supports the *in silico* design strategy demonstrating that the ground state of D66A is similar to the ‘excited’ minor conformational state adopted by the WT, and that the design process effectively swapped the major and minor conformers.

Having established that the ground state of D66A resembles the ‘excited’ state of WT, we acquired CPMG and R1ρ data to further characterise its millisecond dynamics. The raw exchange rates *R*_ex_ for D66A in the 70s loop *R*_*ex*_ were found substantially smaller than those detected for the WT protein, and residues that had large *R*_ex_ values in WT were reduced to zero in D66A (e.g. A66, G74 Fig. 5) indicating the success of the design protocol. Consistent with this, the pattern of specific residues involved with the conformational dynamics was very different (Fig. 5, Table S3) to WT. As with WT, the data could be well explained by fitting to a 2-state model, where the exchange rate was similar to WT (*k* _ex_ ca. 2,000 s^-1^), but the population of the excited state was much smaller, very close to our detection threshold (*p*_B_ ca. 0.5%). Analysis of the excited state chemical shifts shows that this new state is not moving towards a particularly disordered state, nor the ground state of CypA. Instead, since a number of residues in the 80s loop were found to have relatively large *R*_ex_ values in D66A this suggests that the excited state involves rearrangements of the 80s loop, out with the scope of the present simulations that were designed to resolve millisecond timescale motions of the 70s and 100s loops. Overall the key finding is that the NMR data shows that the ‘ground’ conformational state of D66A has the 70s loop in an ‘open’ conformation, which from the chemical shifts, is similar to the ‘excited’ state adopted by the WT protein.

## Discussion

The work demonstrates that our new hybrid method aMD/MSM can accurately and reliably provide a comprehensive description of protein free energy surfaces, including a characterisation of sparsely populated conformational states with millisecond lifetimes. The simulations are hypothesis generating, and are sufficiently accurate to be amenable to rational design strategies whose goal is to adjust the relative populations of the different conformational states (Fig. 3).

It is interesting to compare different ensembles of WT CypA available in the literature. A notable ensemble of WT CypA ^61^ was calculated recently from solution-state NMR data. Our aMD/MSM ensemble is also in good agreement with NOE, J-coupling and RDC solution-state NMR data (Figure S3). However while both ensembles suggest large amplitude motions of the 70s loop, the NMR ensemble does not support the critical role of D66 in a conformational change between open and closed loop conformations, and is also inconsistent with the closed 70s loop conformation observed by X-ray diffraction experiments.

To further test our method, we constructed an ensemble of CypA purely from our aMD trajectories. Unlike the aMD/MSM ensemble, the resulting aMD ensemble predicts that CypA adopts predominantly a 100sopen/70sopen conformation (Fig. S11), a prediction inconsistent with experimental data. Taken together, the aMD/MSM ensemble is the only method that characterises the conformational exchange, revealing the presence of both open and closed conformations of 70s loop in the WT with sufficient precision to provide the means to redesign the conformational landscape.

The resulting conformational states of |CypA also provide insight into function. The open conformational state of the 70s loop has been linked to restriction of the HIV-2 virus. A double mutant (D66N/R69H) of the rhesus macaque homologue TRIMCyp adopts a 70s open conformation, which has been linked to potent HIV-2 restriction.^78^ These observations lead to the interesting speculation that the sparsely populated open 70s conformation in human CypA plays an important and unrealised role in the pathway of HIV-2 infectivity, suggesting stabilisation of this conformation could lead to novel therapeutics.

Approaches have been developed based on using antibodies to stabilise transiently populated conformation states to facilitate ligand discovery efforts ^79–81^. Our results open an exciting opportunity for determining this *in silico* as part of rational drug design endeavours. In the specific case of CypA, current treatments methods have insufficient isoform selectivity^53,54^. We envision that aMD/MSM can be used to selectively target transient states that are differently populated among different protein isoforms, hence paving the way for the rational drug design of is o form selective inhibitors.

There has been substantial discussion on the relationship between millisecond timescale motions in CypA and its catalytic cycle.^82^ Initial measurements suggested that the two could be linked, although this has been disputed by later more detailed measurements ^60,62^. Our results indicate that the opening and closing of the 70s loop is largely independent of the rest of the protein, supporting the notion that its fluctuations are disconnected from the active site.

There remain technical challenges when comparing aMD/MSM simulations to experimental data. To achieve this, it is necessary to calculate chemical shifts from the structural ensembles and combine them appropriately to encapsulate the effects of intermediate exchange in order to accurately capture the effects of millisecond dynamics in NMR data. Chemical shift calculation algorithms such as shiftX2 are trained on static structures and not MD trajectories, and so their accuracy in such applications has not been extensively validated. Nevertheless, the agreement between the predicted and measured CSPs between WT/D66A and H70A/WT are very encouraging. However agreement between *R*_ex_ values calculated for one protein calculated in isolation is less impressive, perhaps suggesting that systematic deficiencies in the chemical shift calculations are removed when one takes the difference between two simulated states. Moreover, when using the expectation values for the CSPs from shiftX2 with the rate constants predicted by the MSM leads to a large overestimation of *R*_ex_ values, on the order of hundreds per second. This suggests that a combination of both the populations and the kinetics together with the estimated chemical shifts are together not in perfect agreement. This can be partially explained by noting that while the CPMG and R1ρ data was acquired at 10 °C, the simulations were performed at 25 °C. These technical challenges aside, it is clear that the main predictions of aMD/MSM are fundamentally correct, and the key conformational ensembles predicted are sufficiently accurate to enable rational design efforts and to generate and test structural hypotheses.

Overall we demonstrate that aMD/MSM can provide atomic-level insights into the thermodynamics and kinetics of independent conformational sub -ensembles that characterise the free energy surface of CypA. The picture is chemically intuitive, and sufficiently accurate for rational design efforts aimed at stabilising particular conformational states over others, even when the lifetimes of the individual states reaches the millisecond timescale. We anticipate that this protocol can greatly facilitate future protein structure-dynamics-function relationships studies and will unlock new opportunities in bioengineering or drug discovery.

## Methods

Molecular dynamic simulations were performed using AMBER16^83^ and GROMACS5.0^84^ software packages and processed using MDTRAJ.^85^ MSMs were built and analysed using PyEMMA 2.3.0.^86^ PALES^87^ and BME^88^ were used to compute RDCs, chemical shifts were calculated with ShiftX2^76^ and MultiEx^89^. NMRPipe^90^ and Sparky3.1^91^ were used to process all spectra. The CCP4 suite^92^ was used for the structure refinement of crystallographic data. Detailed simulation and experimental methods are provided in the Supporting Information.

## Supporting information

Detailed Materials and Methods section, MSM validation procedures, LC-MS analysis for CypA and D66A, additional NMR experiments, CypA assignment, X-ray Refinement statistics, exchange parameters for CypA and D66A, measured *R*_ex_ values. Instructions to download the available datasets. Figures S1-S12. Tables S1-S5.

## Acknowledgements

Authors thank Prof. Beat Vogeli and Prof. Carlo Camilloni for kindly providing sets of NMR observables for Cyp A, and Dr. Xavier Hanoulle and Prof. Guy Lippens for kindly providing the NMR residue assignment of wild type CypA. J. M. was supported by a Royal Society University Research Fellowship. The research leading to these results has received funding from the European Union’s Horizon 2020 research and innovation program under the Marie Sklodowska-Curie grant agreement No. 655667 awarded to J. J-J. and from the European Research Council under the European Seventh Framework Programme (FP7/2007-2013)/ERC grant agreement No. 336289 and from EPSRC (grant no. EP/P011330/1). This project made use of time on the ARCHER UK National Supercomputing Service (http://www.archer.ac.uk) granted via the UK High-End Computing Consortium for Biomolecular Simulation, HECBioSim (http://hecbiosim.ac.uk), supported by EPSRC (grant no. EP/L000253/1). This work was supported by Wellcome Trust Multi-User Equipment grant 101527/Z/13/Z and the Diamond Light Source for beamtime (BAG proposal MX18515). Gratitude is expressed to Dr. Juraj Bella for assistance with experiments at the Edinburgh biomolecular NMR unit.

## Graphical TOC

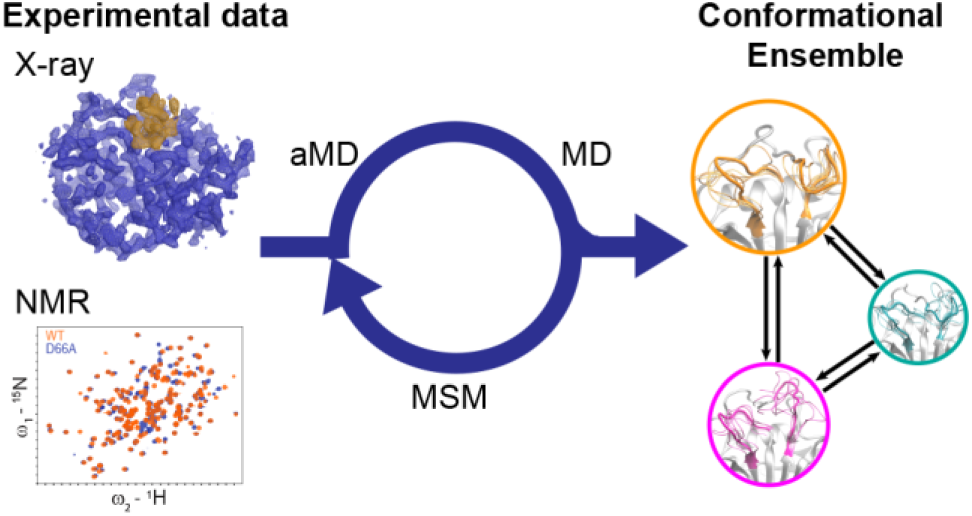

