## Supplementary Information for "Dynamic design: manipulation of millisecond timescale motions on the energy landscape of Cyclophilin A"

**Table of content**


**Materials and Methods: Computational Methods** S4

Accelerated Molecular Dynamics (MD) S4

Molecular Dynamics (MD) S5

MSM Generation S6

MSM Coarse graining S7

Ensemble observables S8

**Materials and Methods: Experimental Methods** S13

Expression Vectors S13

Protein Expression and Purification S13

NMR Methods S14

Isothermal Titration Calorimetry S17

Crystallisation, Data Collection and Structure Determination S18

Synthesis of Ethyl n-[(4-aminobenzyl)carbamoyl] glycinate (**1**) S18

**Supplementary Information References** S19

**Supplementary Figures**

**Figure S1** S25

**Figure S2** S26

**Figure S3** S27

**Figure S4** S29

**Figure S5** S30

**Figure S6** S31

**Figure S7** S32

**Figure S8** S33

**Figure S9** S34

| | |
|---|---|
| **Figure S10** | S35 |
| **Figure S11** | S36 |
| **Supplementary Tables** | |
| **Table S1** | S37 |
| **Table S2** | S44 |
| **Table S3** | S45 |
| **Table S4** | S46 |
| **Available Datasets** | S50 |

**Materials and Methods: Computational Methods**

Accelerated Molecular Dynamics (aMD)

Initial exploration of the conformational landscape of CypA was performed by means of aMD[1,2] using the human Cyclophilin A structure from PDB ID: 1AK4[3] as starting point. Standard MD protein preparation procedures, including removal of co-crystallized proteins and co-solvent molecules and addition of missing hydrogen atoms, were followed. The protein was assigned ff14SB atom types and then solvated on an octahedral TIP3P[4] water box of 12 Å of radius and 1 $Cl^-$ anion was added to neutralize the system. Prior to the aMD runs, the system was minimized combining 4500 steps of steepest descent followed by 7500 steps of conjugate gradient and subsequently heated from 100 K to 298 K in two stages of 125 ps (from 100 K to 200 K and from 200 K to 298 K) at constant volume. Density of the system was then equilibrated by means of a 250 ps long simulation at 298 K in the NPT ensemble. Throughout the heating and equilibration stages SHAKE[5] was applied to all bonds involving hydrogens and a time step 1 fs was employed. Finally, a conventional 100 ns simulation in the NTV ensemble, applying SHAKE to bonds involving hydrogen and using a 2 fs timestep, was performed.

During aMD runs, a positive energy boost is added to the potential energy function, therefore allowing for a simultaneous speed up in the sampling of multiple degrees of freedom without relying on a set of predefined collective variables. The boost potential ($\Delta V(\boldsymbol{r})$) is only applied when the average potential energy of the system ($V(\boldsymbol{r})$) falls below a certain threshold ($E_P$) and, therefore, the potential used in aMD simulations takes the form of eq 1.

$$V*(r) = \begin{cases} V(\boldsymbol{r}) & V(\boldsymbol{r}) > E_P \\ V(\boldsymbol{r}) + \Delta V(\boldsymbol{r}) & V(\boldsymbol{r}) < E_P \end{cases} \qquad (1)$$

The current implementation of aMD in PMEMD[2] allows applying a dual boost approach where an energy boost is applied to all heavy atoms in the system and an additional boost is applied solely to the torsional degrees of freedom, hence the boost potential is given by eq 2.

$$\Delta V(\boldsymbol{r}) = \frac{(E_P - V(\boldsymbol{r}))^2}{(\alpha_P + E_P - V(\boldsymbol{r}))} + \frac{(E_D - V_D(\boldsymbol{r}))^2}{(\alpha_D + E_D - V_D(\boldsymbol{r}))} \qquad (2)$$

where $E_P$ and $E_D$ are the reference potential and torsional energies respectively; $V_D$ is the dihedral component of the potential energy and $\alpha_P$ and $\alpha_D$ serve to control the intensity of the boost applied to each term. Following guidelines of Pierce and co-workers[2], and the information obtained from the preliminary MD, a value of -81941 kcal/mol was used for $E_P$, while $E_D$ was set to 2535 kcal/mol. After trial runs (data not shown), values of 2267 and 66 were used for $\alpha_P$ and $\alpha_D$ respectively. One common limitation of aMD is that a too intense energy boost may push the sampling towards high-energy regions of the Potential Energy Surface (PES), effectively triggering protein unfolding. To surpass this limitation harmonic restraints, using a force constant of 15 kcal mol$^{-1}$ A$^{-2}$, were applied to the backbone atoms of residues that display α-helix or β-strand secondary structure in the crystallographic X-ray structure with PDB code 1AK4. At this stage, protein preparation was performed with tLeap and simulations were carried out employing the GPU accelerated version of PMEMD[6], both from the AMBER16[7] software package.

Molecular Dynamics (MD)

Structures sampled from the aMD trajectory were used as initial structures to seed sets of MD trajectories. Virtual sites were used in order to allow for a time step of 4 fs during MD simulations[8,9]. FF14SB atom types were assigned to the protein and each structure was then solvated in a 12 Å of radius dodecahedral water box and 1 $Cl^-$ anion was added to neutralize the system. Each system, comprising ca. 30000 atoms, was minimized combining 4500 iterations of steepest descent with 5000 iterations of conjugate gradient. Subsequently systems were heated from 0 K to 250 K in 150 ps in the canonical ensemble using a time step of 0.5 fs, and from 250 to 298 K in 300 ps using a timestep of 1 fs. Prior to the production runs each system was equilibrated for 300 ps at 298 K and 1 bar of pressure using 2 fs timestep, before switching to a 4 fs timestep for 10 ns under the same temperature and pressure conditions. Each system was then used for production runs consisting of 150 ns long trajectories in the NTP ensemble using a 4 fs timestep. MD simulations were set up and conducted using the appropriate modules of the GROMACS5.0[10,11] software package, using LINCS[12] algorithm truncating the constraint coupling matrix at the sixth order.

MSM Generation

MSMs were built from the set of equilibrium MD trajectories using the pyEMMA 2.3.0 software package.[13] In order to achieve a successful dimensionality reduction for the MSM building movements of the 70s loop (spanning from residue 65 to 77) were accurately captured by the root mean square deviation of the loops Cα atoms with respect to their position in the X-ray structure 1AK4. However, the Cα RMSD of the 100's loop between (spanning from Met100 to Ser110) failed to describe the flapping movement found by visual inspection of the trajectories, because significantly different snapshots were mapped to the same RMSD values. In order to accurately describe this movement the distance between the centre of mass (COM) of the 100s loop and the α-helix defined by residues Pro30 to Thr41 was simultaneously monitored. Subsequently, a negative value was assigned to those values of RMSD where such distance was shorter than in the X-ray structure (14 Å). The dimensionality of the MD trajectories was reduced by projecting onto these two RMSD based collective variables. K-means clustering using 100 clusters was used to obtain a microstate definition for the MSM construction. Different lagtimes were used to compute implied timescales of the dominant eigenvalues according to eq 3:

$$t_i = -\frac{\tau}{ln\,\lambda_i(\tau)} \qquad (3)$$

The resulting implied timescale ($t_i$) plots are shown in Figure S1. A Bayesian MSM was built at each lagtime (τ) in order to obtain an error estimate for the obtained timescales. Based on the implied timescales a lagtime of 40 ns was chosen for the MSM construction, with default parameters provided by PyEMMA.[14]

MSMs construction for the mutants was carried out using the same features for dimensionality reduction, and initial snapshots were obtained by re-using those from the CypA seeds, after mutation of appropriate residues. To facilitate comparative analyses the MSMs for CypA and the mutants were built on a common CV space and set of cluster centres obtained from clustering a combined set of 2000 MD trajectories (1200 for CypA and 400 for each mutant) amounting for a cumulative sampling time of ca. 260 μs.

MSM Coarse graining

Based on the eigenvalue structure of the transition matrix of the MSM, five macro-states were deemed useful for describing the slow dynamics of the system. The 100 k-means derived model was therefore coarse grained into 5 metastable conformations using a hidden Bayesian Markov Model.[15,16] This grouping successfully distinguished between ordered, disordered and intermediate states of the 70s-loop, but clustered together structures on both ends of the 100s loop CV range and did not allow for a direct comparison between CypA and D66A, due to the narrow separation of timescales in the mutant protein. A refinement of the coarse grained model was therefore achieved according to the following structural criteria: microstates with a value of the 70s-loop CV below 1.5 Å and with a value of the 100s-loop

CV below 4.5 Å were assigned to the ground state (closed/open, orange). Microstates within the same cutoff of the 70s-loop CV but with values of the 100s-loop above 4.5 Å were assigned to the closed/closed metastable state (red). Microstates with values of the 70s-loop CV between 1.5 Å and below 4.0 Å were assigned to the intermediate macro-state (teal). Microstates with a 70s-loop CV value above 4.0 Å and a 100s-loop CV value < 4.5 Å were assigned to the open/open state (magenta) while microstates with similar values of the 70s-loop CV but extreme values of the 100s-loop CV (>4.5 Å) were assigned to the open/closed (blue) macro-state. This structural assignment was found to be the one that best reproduced the hidden Bayesian Markov Model while allowing for direct comparison between CypA and mutant proteins. The robustness of the calculated assignment was asserted by recomputing the predicted populations of each macro-state whilst varying fractions of the total MD dataset (Figure S2). Mean first passage times (MFPTs) between pairs of non-adjacent macro-states states were computed as a weighted average of MFPTs over all pairs of microstates. MFPT between adjacent macro-states were computed between the two most populated microstates of each microstate.

Ensemble observables

*Previously published structural ensembles:* Comparison with experimental structural ensembles of CypA was performed by obtaining the NMR ensemble with PDB ID: 2N0T and the X-ray structure with PDB ID: 1AK4. The calculated ensemble for CypA was analysed by including all macrostates, or just the dominant macrostate (orange). This was done to determine how sensitive the observables are to the different loop conformations present in the ensemble.

*NOEs and eNOEs*: The distance between each of the 1253 atom pairs involved in eNOE signals and 2143 involved in NOE signals was calculated for each structure in the ensemble with MDTRAJ,[17] using CYANA definitions[18] for pseudo atoms representing nondistinguishable hydrogen atoms. Subsequently, a histogram weighted by the probability of each microstate of the MSM was built and the corresponding NOE signal between hydrogens $i$ and $j$ for each histogram was calculated as in equation (4):

$$r_{i,j}^{avg} = \langle r_{i,j}^{-6} \rangle^{\frac{-1}{6}} \quad (4)$$

Each predicted signal was compared with the experimental value and for the eNOE signals it was considered fulfilled if the predicted value was within lower and upper limits ± a tolerance value of 0.5 Å. Standard NOE signals were considered fulfilled if the predicted value was below the upper limit derived from experiment. Errors were estimated by 1000 cycles of bootstrapping of the histograms from individual trajectories to generate ensemble histograms. The results are reported in Figure S3a.

*Coupling Constants ($^3J$)*: Backbone ϕ/ψ dihedral angles were calculated with MDTRAJ for all the structures in the ensemble. Karplus[19,20] equations were used to obtain values for each type of $^3$J and a histogram weighted by the probability of each microstate was subsequently built. Error estimation was performed as for the prediction of NOE and eNOE. The results are reported in Figure S3b.

*Residual Dipolar Couplings (RDCs)*: A representative set of 1000 snapshots gathered according to microstate probabilities of the Markov model was used to compute an average structure after removing translation and rotation movements. Subsequently, PALES[21] was used to calculate an alignment tensor that provided the best fit of experimental residual couplings for residues in the β-sheet and α-helix regions of CypA to this average structure using singular value decomposition.[22] A subset of 5000 structures of the conformational ensemble was used to compute average RDCs, and the corresponding Q value was calculated. For the MD ensemble an average weighted by the probability of each microstate was calculated, while for the NMR ensemble available in the literature an unweighted average over all structures was used. Errors on the average Q values were estimated by bootstrapping.

The Q factor considering only residues in helices and sheets is 0.23 indicating a good quality description of the overall fold by the MD simulations. This increases to 0.31 when flexible loops are also considered in the calculations. These figures are similar to those obtained for the X-ray structure of CypA. The poorer description of RDCs for residues in the flexible loops of CypA is not surprising considering the large amplitudes of backbone fluctu ations. Zweckstetter, M. & Bax[23] have shown that a Q factor of ca. 0.35 corresponds to an average error in the alignment angle of 10 degrees. In addition the RDC back-calculation procedure from the MD ensemble used here is only rigorously valid for conformers in fast exchange, whereas the experimental data was measured for several loops residues that are in intermediate exchange. It is difficult to quantify the errors introduced by this approximation, but this may explain at least in part why the Q-factors for the flexible loops are larger than for the rigid regions.

To account for model errors caused by incomplete sampling and limited accuracy of the potential energy function used we also used the Bayesian Maximum Entropy reweighing tool BME to improve agreement with experimental data for the RDCs.[24] The procedure causes small changes in the populations of the 5 MSM macrostates (blue 0.08 —> 0.10; magenta 0.18 —> 0.09; orange 0.41 —.> 0.54; red 0.28 —> 0.24; teal 0.05 —> 0.01). These changes do not qualitatively affect the ranking of the macro states by populations, though overall the 70s closed states become more populated (ca. 70 to 90%). This leads to a Q-factor of the rigid regions of CypA is 0.20 and including the flexible loops the Q-factor is 0.21, thus the flexible loops are as well described as the rigid regions in the reweighted ensemble. Q-factor values of ca. 0.20 indicate that the structural description of the protein is of good quality.[25]

By comparison the Chi et al. NMR ensemble has a Q-factor of 0.36 for the rigid CypA regions and this increases to 0.37 when flexible loops are also considered. We also attempted to improve agreement against experimental data by reweighing the Chi et al ensemble with the software BME. In this case the best reweighed ensemble has a Q-factor of 0.31 (rigid regions only) and 0.30 (considering flexible loops as well). Camilloni et al. reported a global Q factor of 0.31 for a separate MD ensemble of CypA refined against chemical shifts.[26] Thus overall the present MD ensemble appears to more accurately account for the RDCs of the flexible loops than the other ensembles. A summary of these results is reported in Figure S3c.

*Chemical shifts:* For both CypA and D66A chemical shifts of backbone $^{15}N$ and $^{1}H$ nuclei for each microstate of the MSMs were estimated by sampling 100 snapshots from each microstate, and using ShiftX2[27] to calculate the average chemical shift value of each residue. The MSM transition probabilities were converted into a rate matrix and input to the software MultiEx together with the average chemical shift. MultiEx is an *N* site generalization of the *cpmg_compare* software that was previously written to simulate numerically CPMG for two site chemical exchange.[28] MultiEx takes as input chemical shift values for an *N* state system and a rate matrix and provides simulated HSQC spectra that were used to produce the CSP plots. The chemical shifts of the NMR ensemble were computed assuming fast exchange between all structures in the ensemble. A summary of the overall results for WT CypA is given in Figure S3d.

Overall (Fig S3) the results for the above observables between the one macrostate and 5 macrostate ensemble are similar, indicating that these cannot reliably validate the existence of minor states in CypA and therefore that additional CPMG/R1p experiments are necessary to validate the proposed ensemble.

*HetNOE values:* A 2.5 μs MD simulation of CypA was initiated from a snapshots sampled from a well populated cluster of the MSMs of WT and D66A respectively. Snapshots were saved every 5 ps. The simulations were split in two separate trajectories of 1.25 μs each and analysed separately to estimate uncertainties in the calculated HetNOE values according to a protocol based on work from Robustelli et al.[29] Firstly N-H bond vector autocorrelation functions $C_{\mathrm{int}}(t)$ were computed with eq 5 by aligning all snapshots to a reference structure to remove global tumbling motions with a maximum lag time $\tau$ of 100 ns.

$$C_{int}(t) = \langle P_2\,[\mu(t) \cdot \mu(t+\tau)]\rangle \qquad (5)$$

where $P_2[x]$ is the second Legendre polynomial of x, μ is the relevant bond vector scaled to unit magnitude, and angular brackets indicate averaging over time $t$. Ill-behaved $C_{\mathrm{int}}(t)$ curves that showed no sign of converging towards a plateau region were discarded from further analyses. Next total autocorrelation functions were obtained by multiplying the $C_{\mathrm{int}}(t)$ curves a global tumbling function $C_{\mathrm{glob}}(t)$

$$C_{tot}(t) = C_{int}(t)C_{glob}(t) \qquad (6)$$

where $C_{\mathrm{glob}}(t) = \exp(-t/\ t_g)$ with $t_g$ taken as the global tumbling time of WT and D66A, as determined from NMR relaxation measurements (9.1 ns and 9.2 ns respectively). Next orientational spectral density functions were calculated by discrete Fourier transform of the resulting simulated autocorrelation functions using eq 7

$$J(\omega) = \frac{2}{5}\mathrm{Re}\left[\int_{-\infty}^{+\infty} C_{tot}(t)e^{-i\omega t}dt\right] \qquad (7)$$

Steady-state heteronuclear $^{1}H$-$^{15}N$ NOE values were determined with eq 8 that assumes only contribution from dipolar interactions:

$$HetNOE = 1 + \frac{\gamma_H}{\gamma_N}\frac{[6J(\omega_H + w_N) - J(\omega_H - \omega_N)]}{J(\omega_H - \omega_N) + 3J(\omega_N) + 6J(\omega_H + \omega_N)} \qquad (8)$$

where $\gamma_H$ and $\gamma_N$ are the gyromagnetic ratios of hydrogen and nitrogen, and $\omega_H$ and $\omega_N$ are the proton and nitrogen resonance frequencies on a 600 MHz spectrometer for compatibility with the measurements carried out in this work. Uncertainties are reported as the standard error of the mean of the *HetNOE* values obtained from analysis of two separate 1.25 μs trajectories.

**Materials and Methods: Experimental Methods**

Expression Vectors

A wild-type $PPIA_{(1-165)}$ gene inserted downstream of an N-terminal hexa-histidine tag with one TEV protease cleavage site in pDEST™ 14 expression vector[30] was provided by the Edinburgh Protein Production Facility (EPPF). Site directed PPIA-D66A mutant gene construct was synthesized by GeneArt (ThermoFisher Scientific) and was subsequently subcloned into a modified pET-15b vector with N-terminal hexa-histidine tag and a thrombin protease cleavage site.

Protein Expression and Purification

Expression and purification CypA was achieved as previously described.[30] The D66A mutant clone construct was transformed into BL21(DE3)pLysS host cells and kept for incubation at 37°C in LB medium. Overnight grown bacterial cultures were induced with 1.0mM isopropyl-β-D-1-thiogalactopyranoside (IPTG) and then further incubated at 30°C for 4-6 hrs. Cells were harvested and lysed in phosphate buffer saline (P.B.S), pH 7.4 as lysis buffer using Constant Cell Systems Disruptor (1.1kW TS Benchtop) set at 25kpsi. The lysed fraction was centrifuged, and directly applied on a $Ni^{2+}$-affinity (GE Healthcare) column. Elution of fusion histidine tagged D66A was achieved with a step wise gradient of 20mM phosphate, 150mM NaCl, 500mM Imidazole, pH 7.4 buffer. Eluted fractions were concentrated and fusion protein sample was further kept for cleavage with thrombin protease at a 1:100 (w:w) protease:protein ratio at room temperature. The cleaved protein samples was further applied to a $Ni^{2+}$-affinity column and flow–thru protein fractions were collected with a loading buffer (20mM phosphate, 150mM NaCl, 30mM Imidazole, pH 7.4). The final protein purification was achieved by passing concentrated samples through a 16/60 Superdex75 size-exclusion chromatography (AKTA™ pure, GE Healthcare) pre-equilibrated with 50mM phosphate, 1mM tris-(2-carboxyethyl) phosphine (TCEP), pH 6.5. $^{13}C$,$^{15}N$ uniformly enriched D66A protein was prepared by following the protocol described above using M9 minimal culture medium with ($^{15}N$) $NH_4Cl$ as nitrogen source and D-($^{13}C_6$) glucose as carbon source as previously described.[31] The purity of unlabelled and labelled CypA and D66A samples was initially confirmed by SDS-PAGE and further by ESI-TOF mass spectrometric analysis. LC-MS profiles of unlabelled CypA and 15N-labelled D66A are shown in Fig S6 and Fig. S7 respectively.

NMR Methods

3D backbone NMR measurements were performed on a Bruker Avance 800MHz spectrometer with a cryogenic TCI probe at 25°C. NMR sample consisted of 1.0mM of $^{13}C$-,$^{15}N$-labelled D66A in 50mM phosphate, 1mM TCEP, (pH 6.5) and 10% $D_2O$ (v/v). Backbone resonance assignment was achieved using the complementary set of experiments of HNCA and HN(CO)CA, HNCO and HN(CA)CO, CBCANH and CBCA(CO)NH[32] acquired on uniformly labelled sample. Partial side chain resonances were assigned from combined information content of ct-$^{13}C$ HSQC and HBHA(CO)NH.[32] All spectra were processed with the NMRPipe[33] and analysed using Sparky 3.1[34] respectively. For automated sequence-specific NMR assignment, PINE-SPARKY[35] a probabilistic algorithm program was used. Secondary structure and backbone dihedral angles were predicted using TALOS+[36] using D66A chemical shifts assignment as input. A total 143 backbone residues out of 165 amino acids were assigned on D66A, excluding 6 prolines that could not be detected in $^{1}H$-

$^{15}$N HSQC spectra. Missing resonances for His70, Asn71, Thr73, Lys76 and Glu81 in D66A HSQC spectrum were attributed to the effect of the mutant on the backbone dynamics of the 70s-loop region. Chemical shifts for D66A have been deposited in BMRB (BMRB entry 12023).

*Chemical Shift Changes.*

The overall chemical shift changes (Δω) were determined using equation (9):

$$\Delta\omega = \sqrt{\left(5\Delta\,^{1}H\right)^{2} + \left(\Delta\,^{15}N\right)^{2}} \qquad (9)$$

Here, $\Delta^{1}$H and $\Delta^{15}$N are the changes in the proton and nitrogen dimensions, respectively. Figure S8 highlights some significant CSPs.

*$^{15}$N NMR Relaxation Measurements*

Backbone T1 and T2 relaxation measurements were carried out using with 1.0-1.3mM $^{15}$N-labeled CypA and D66A using previously described pulse sequences[37] in a interleaved manner at two different field strengths of 500 and 600MHz at 25°C. Eight $T_1$ relaxation times were sampled from 100ms to 1000ms with a replicate point at 300 ms for error analysis. Triplicate set of {$^{1}$H}-$^{15}$N heteronuclear NOE (hetNOE) relaxation measurements were recorded at both field strengths of 500 and 600MHz using water flip-back NOE pulse sequence in an interleaved fashion with 5 s relaxation delay to compute the uncertainties within the measured hetNOE values. $T_2$ relaxations delays at eight different time points ranging from 20 to 150 ms with a replicate point at 20 ms were acquired. 2D datasets were processed using NMRpipe[33] and arrayed HSQC spectral cross peaks signal intensities as a function of variable time delays were fitted to single exponential decay using FuDa[38] to determine the $R_1$ and $R_2$ relaxation rates. HetNOE values were calculated by taking the ratio of peak intensities according to the equation $I_{on}/I_{off}$ from experiments performed with and without 1H pre-saturation. In total, 142 cross peaks for CypA and 149 cross peaks for D66A were used for analysis.

*Model-free Analysis*

Model-free analysis was performed using $R_1$, $R_2$ and hetNOE values for CypA and D66A with relax software. Residues identified as exchanging on the microsecond-millisecond time scale were excluded from analysis in determining the overall diffusion tensor. The globally fitted correlation times $\tau_c$ for both proteins are determined to be similar: 9.1 ns and 9.20 ns for CypA and D66A respectively. However the analysis suggests that effective correlation times for internal motions of a large number of residues approaches that of the global tumbling time, and thus low values of generalized-order parameters may not reliably indicate fast motions (Figure S9).

*15N Backbone Relaxation Dispersion Experiments*

Relaxation compensated $^{15}$N backbone Carr-Purcell-Meiboom-Gill (CPMG) experiments[39] were carried out on 1.0-1.3mM $^{15}$N-labeled wild-type CypA and D66A CypA variant protein samples. Data were collected at 10°C using multiple magnetic field strengths of 500 and 600MHz. The constant time relaxation delay was set to 32ms. $^{15}$N-relaxation dispersion profiles were generated from the cross-peak intensities obtained from the $^1$H-$^{15}$N 2D datasets extracted for 14 values of the CPMG field strength, varying from 67 to 1000 Hz with two values of $v_{cpmg}$ datasets acquired for error estimation. Peak intensities as a function of CPMG frequency of $v_{cpmg}$ were quantified using FuDa.[38] Individual residue relaxation dispersion curves were evaluated for statistically significant dispersions for two-site or three site exchange model versus no exchange model (flat horizontal line). Residues that displayed appreciable exchange contributions were identified and clustered for global analysis using the numerical simulation software CATIA[40], which fits $k_{ex}$ and $p_b$ globally and a $|\Delta\omega|$ for each exchanging residue. $R_{Nz}$ and $R_{HzNz}$ rates were measured for the proteins and used as inputs for CATIA to determine the differential relaxation rates of in-phase and anti-phase magnetization as well as the peak positions so that the effects of off-resonance pulses are fully accounted for. Off-resonance R1ρ relaxation dispersion experiments on $^{15}$N were recorded at 600MHz field strength at 10°C using the pulse sequence described by Korzhnev et. al.[41] (Fig S7) with offsets $\Omega_0/2\pi$ ranging from -2.0 to 2.0 kHz, $\Omega_{ref}$ being the centre of the $^{15}$N-spectrum (116.5 ppm), spin-lock field strengths ranging from 431 to 1649 Hz and offsets ranging from 0 to 10 000 Hz. See also Figure S10.

Isothermal Titration Calorimetry

Calorimetric experiments were carried out using an Auto-iTC200 micro-calorimeter (Malvern Instruments) at 25 °C. The titration experiments between compound i.e Ethyl n-[(4-aminobenzyl)carbamoyl]glycinate and CypA and D66A protein samples were performed in 50mM phosphate, 1 mM Tris-(2-carboxyethyl) phosphine (TCEP) pH-6.5 buffer. In each experiment, the 200 µl sample cell was filled with 100 µM (CypA) or 200 µM (D66A) protein sample, and the 40 µl injection syringe with 2.0 mM or 5.0 mM compound solution. The titrations were performed with an initial 0.4 µl injection, followed by 19 injections of 2.0 µl at 240 s intervals, while stirring the solution at 750 rpm. Control experiments were also performed with compound titrated to carefully matched experimental buffer. Raw datasets were integrated and baseline corrected using NITPIC.[42] Binding constant, $K_d$ values reported for compound versus CypA variants are results of titration experiments fitted with “ A + B <-> AB’ hetero-association model using the non-linear least square routine in-built within ITCsys and final single-site binding model results plotted with GUSSI.[42]

Crystallisation, Data Collection and Structure Determination

N-terminal hexa-histidine tag was removed from the purified D66A protein by incubation with Thrombin protease for 3 hours at room temperature. The protein was re-purified and concentrated to 25 mg ml$^{-1}$ in X-ray buffer (20 mM Tris pH 7.4, 100 mM NaCl, 1 mM DTT). A stock solution of the reference compound at a concentration of 50 mM was prepared in X-ray buffer plus 25% v/v DMSO. The stock solution was mixed with protein solution at a 1:4 v:v ratio (ligand:protein) and equilibrated in ice for 30 minutes. The ammonium sulfate suite (QIAGEN) was then used to test the crystallisation of the complex. Crystals suitable for X-ray diffraction were grown by vapour diffusion using the sitting drop method at 17 $^{o}$C. Drop consisted from 1:1 ratio of protein-ligand complex and well buffer and crystals were observed after one day in different well positions, including wells B7, D2, D8 and H5. Prior diffraction, crystals were cryoprotected by brief immersion in oil (1250 cSt from Sigma) and flash frozen in liquid nitrogen. X-ray diffraction data were collected at 100K at the Diamond Light Source synchrotron-radiation facility in Oxfordshire, England. Data were processed with MOSFL [43] and AIMLESS [44]. Molecular replacement was used to solve the structure of D66A in complex with the reference ligand using PDB ID 4N1M. Structure refinement was performed using the Crystallographic Object-Oriented Toolkit (Coot) [45] and REMFAC5 software [46] part of the CCP4 suite [47].

Synthesis of Ethyl n-[(4-aminobenzyl)carbamoyl] glycinate (**1**)

Ligand **1** was synthesized applying a previously reported procedure.[48] Spectroscopic data confirmed the structure. The procedure is reported below:

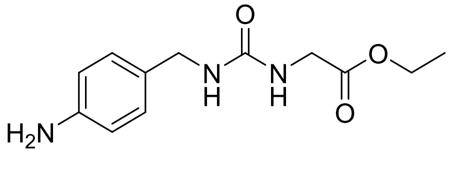


**1**

Ethyl isocyanatoacetate (0.4 mL, 3.68 mmol) was dissolved in DMF (15 mL). 4-Aminobenzylamine (0.4 mL, 3.68 mmol) was added and the reaction mixture was stirred at room temperature for 2 h. After this time, diethyl ether (50 mL) was added and the resultant solid was filtered off, yielding the desired product. White solid 900 mg (97% yield). $^{1}$H NMR (500 MHz, DMSO-$d_6$) δ 6.94 – 6.88 (m, 2H), 6.53 – 6.47 (m, 2H), 6.39 (t, $J$ = 5.8 Hz, 1H), 6.15 (t, $J$ = 6.1 Hz, 1H), 4.92 (s, 2H), 4.09 (q, $J$ = 7.1 Hz, 2H), 4.02 (d, $J$ = 5.7 Hz, 2H), 3.77 (d, $J$ = 6.0 Hz, 2H), 1.20 (t, $J$ = 7.1 Hz, 3H). $^{13}$C NMR (126 MHz, DMSO-$d_6$) δ 171.67, 158.30, 147.85, 128.51 (2C), 127.80, 114.13 (2C), 60.61, 43.20, 42.02, 14.59.

**Supplementary Information References**

6. Salomon-Ferrer, R., Goetz, A. W., Poole, D., Le Grand, S. & Walker, R. C. Routine microsecond molecular dynamics simulations with AMBER on GPUs. 2. Explicit

solvent particle mesh ewald. *J. Chem. Theory Comput.* **9**, 3878–3888 (2013).

15. Noé, F., Wu, H., Prinz, J. H. & Plattner, N. Projected and hidden Markov models for calculating kinetics and metastable states of complex molecules. *J. Chem. Phys.* **139**,

184114 (2013).

24. Bottaro, S., Bengtsen, T. & Lindorff-Larsen, K. Integrating Molecular Simulation and Experimental Data: A Bayesian/Maximum Entropy Reweighting Approach. *bioRxiv*

(2018). doi:10.1101/457952

33. Delaglio, F. *et al*. NMRPipe: A multidimensional spectral processing system based on

UNIX pipes. *J. Biomol. NMR* **6**, 277–293 (1995).

41. Korzhnev, D. M., Skrynnikov, N. R., Millet, O., Torchia, D. A. & Kay, L. E. An NMR experiment for the accurate measurement of heteronuclear spin-lock relaxation rates. *J.*

*Am. Chem. Soc.* **124**, 10743–10753 (2002).

**Supplementary Figures**

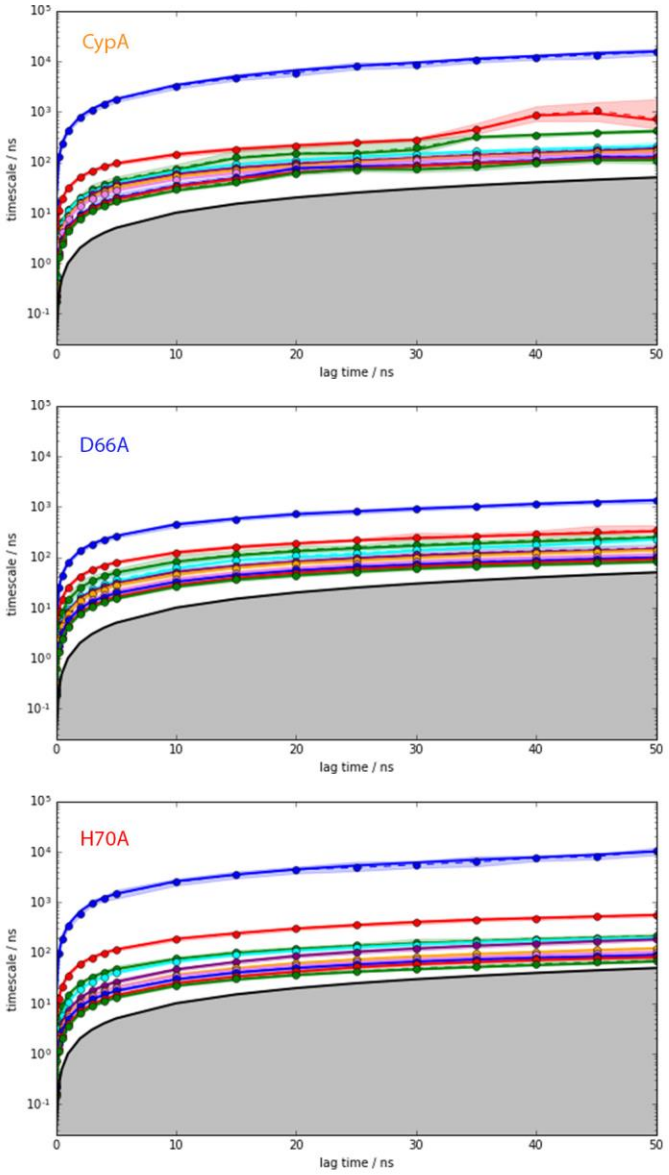


**Figure S1:** Implied timescales of the predicted MSM as a function of lagtime. Shaded region represents the estimated statistical errors.

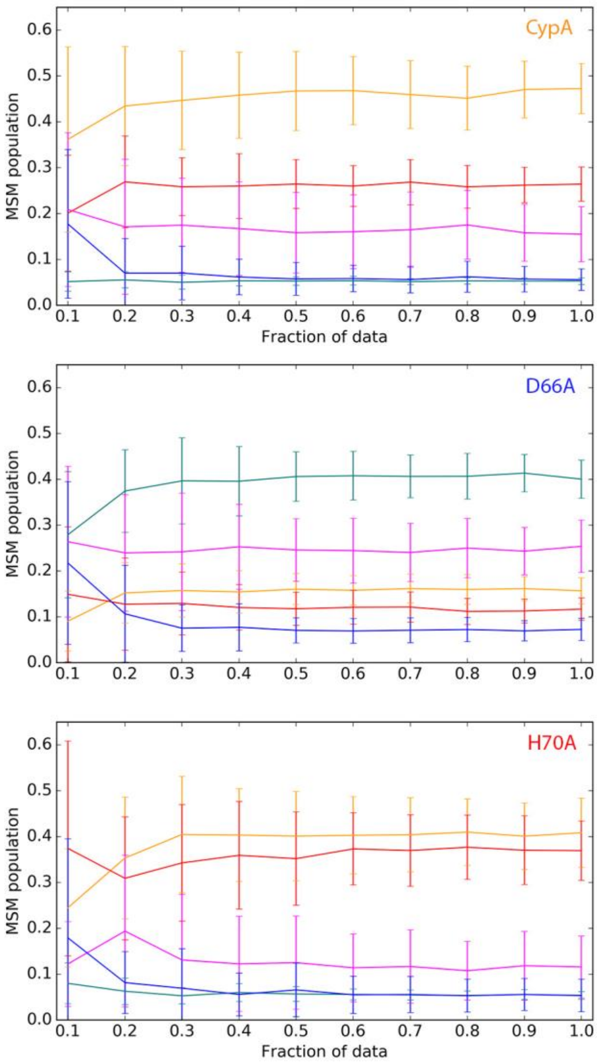


**Figure S2:** Predicted population of each MSM macrostate as a function of the fraction of the total dataset used to build the model. Error bars were obtained by bootstrapping.

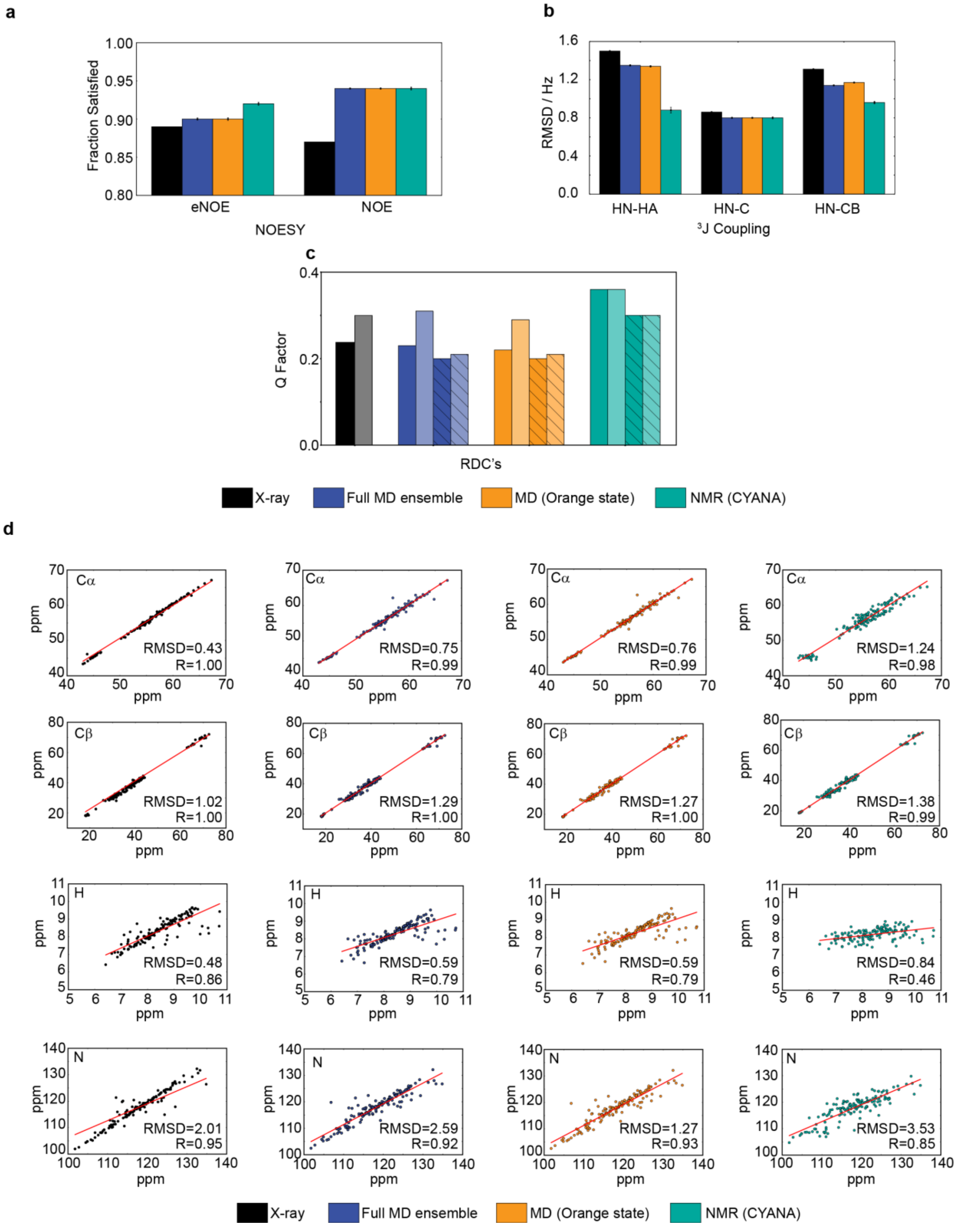


**Figure S3.** Calculated observables for the X-ray structure, MD and NMR ensembles of CypA. **a NOEs**. The MD ensembles reproduce the vast majority of eNOE and NOE-derived protein proton distances. The NMR ensemble was refined against this set of NOEs but shows almost identical accuracy. The X-ray structure is the least consistent with the NOE data. **b J-couplings**. The NMR ensemble is more accurate at reproducing HN-HA and HN-HN couplings and of similar accuracy to the MD ensembles for HN-C couplings. This may be because the ensemble was refined against this set of observables. The X-ray structure is the least consistent with the J-couplings data. **c RDCs**. Dark bars correspond to secondary structure regions, whereas light bars also include the flexible loops. Shaded bars correspond to results obtained with reweighting. See SI text for a detailed discussion. The NMR ensemble is less accurate than the MD ensembles at reproducing RDCs. This may be because the NMR ensemble was not refined against this particular set of RDCs that come from a separate study.[26] **d** Correlation plot of experimental and predicted values of backbone N, H, $C_\alpha$ and $C_\beta$ chemical shifts. The best results are obtained using the crystal structure of CypA, but this may be fortuitous as CypA was included in the training set of crystal structures used to calibrate ShiftX2.[27,49]

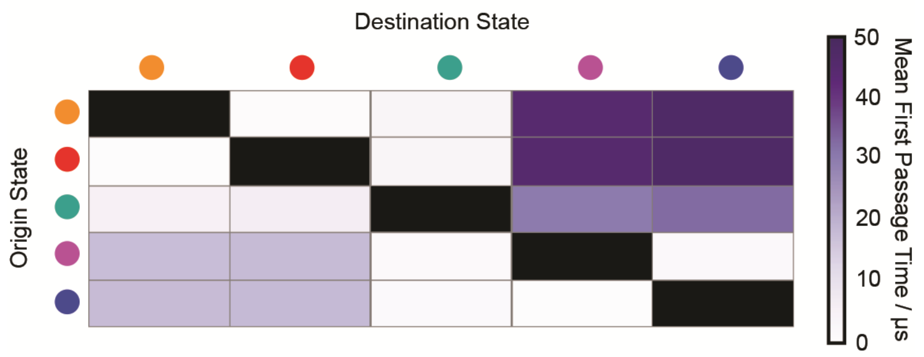


**Figure S4.** Mean First Passage Time among CypA MSM macrostates. Transitions involving the 70s loop disordering is approximately one order of magnitude slower than transitions involving 100s loop flapping.

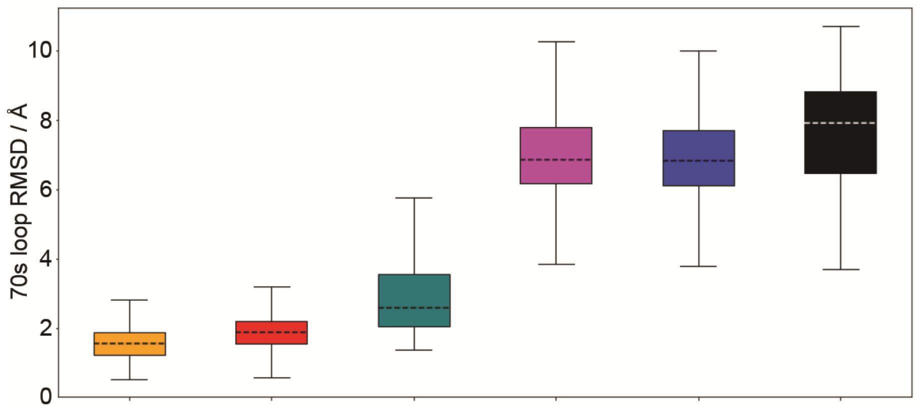


**Figure S5.** Per state boxplot of RMSD values of the 70s loop with respect to reference X-ray structure 1AK4. Each color identifies a macrostate as described in the main text and the 2N0T NMR bundle is shown in black.

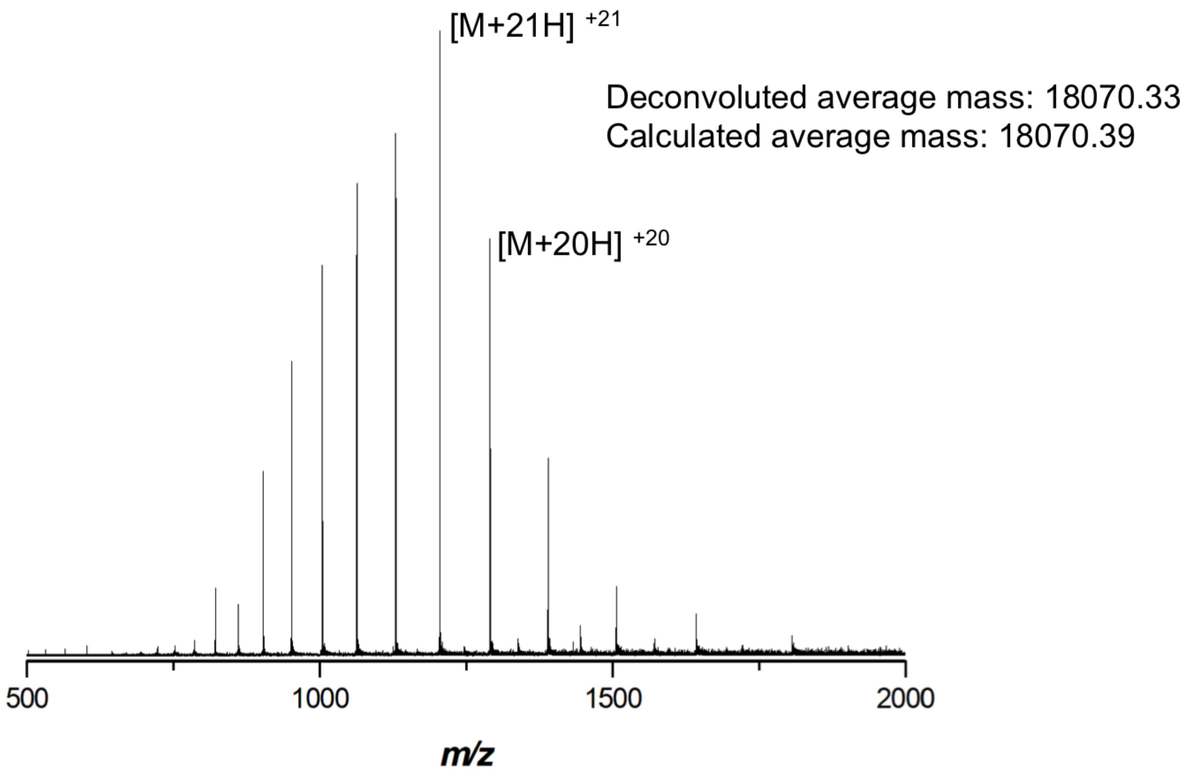


**Figure S6.** Positive mode mass spectrum acquired by LC-MS of wt-CypA following cleavage. A charge state distribution can be seen, and the $[M+21H]^{+21}$ and $[M+20H]^{+20}$ ions have been highlighted. The deconvoluted average mass was calculated to be 18070.33 Da, in agreement with the calculated theoretical average mass of 18070.39 Da.

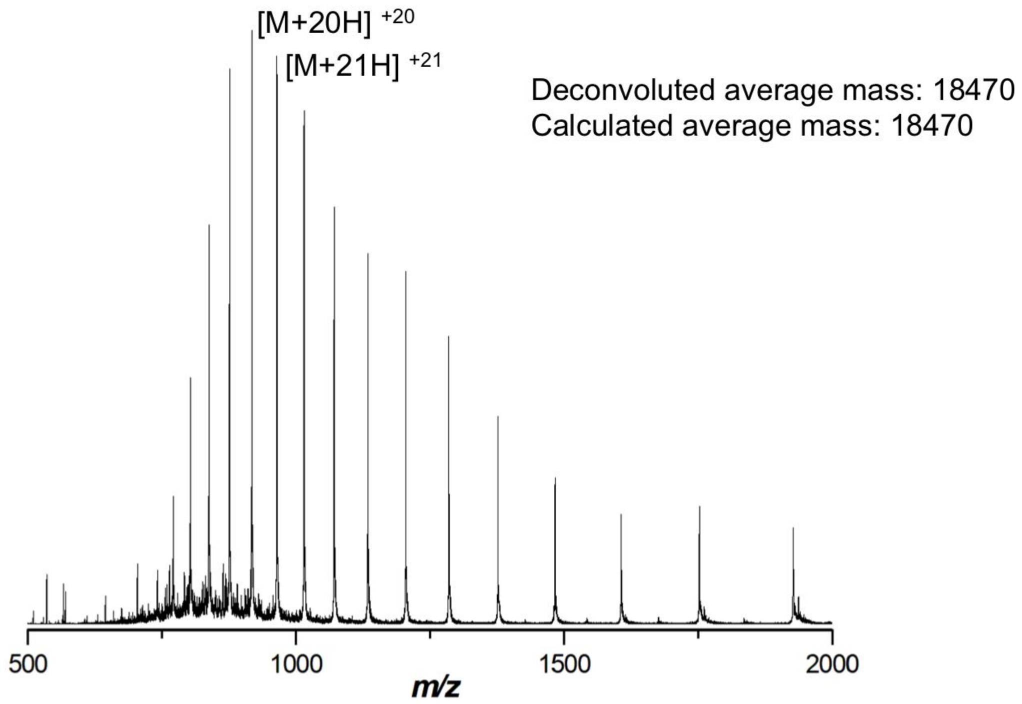


**Figure S7.** Positive mode mass spectrum acquired by LC-MS of $^{15}$N labelled D66A CypA following cleavage. A charge state distribution can be seen, and the $[M+21H]^{+21}$ and $[M+20H]^{+20}$ ions have been highlighted. The deconvoluted average mass was calculated to be 18470 Da, in agreement with the calculated theoretical average mass of 18470 Da.

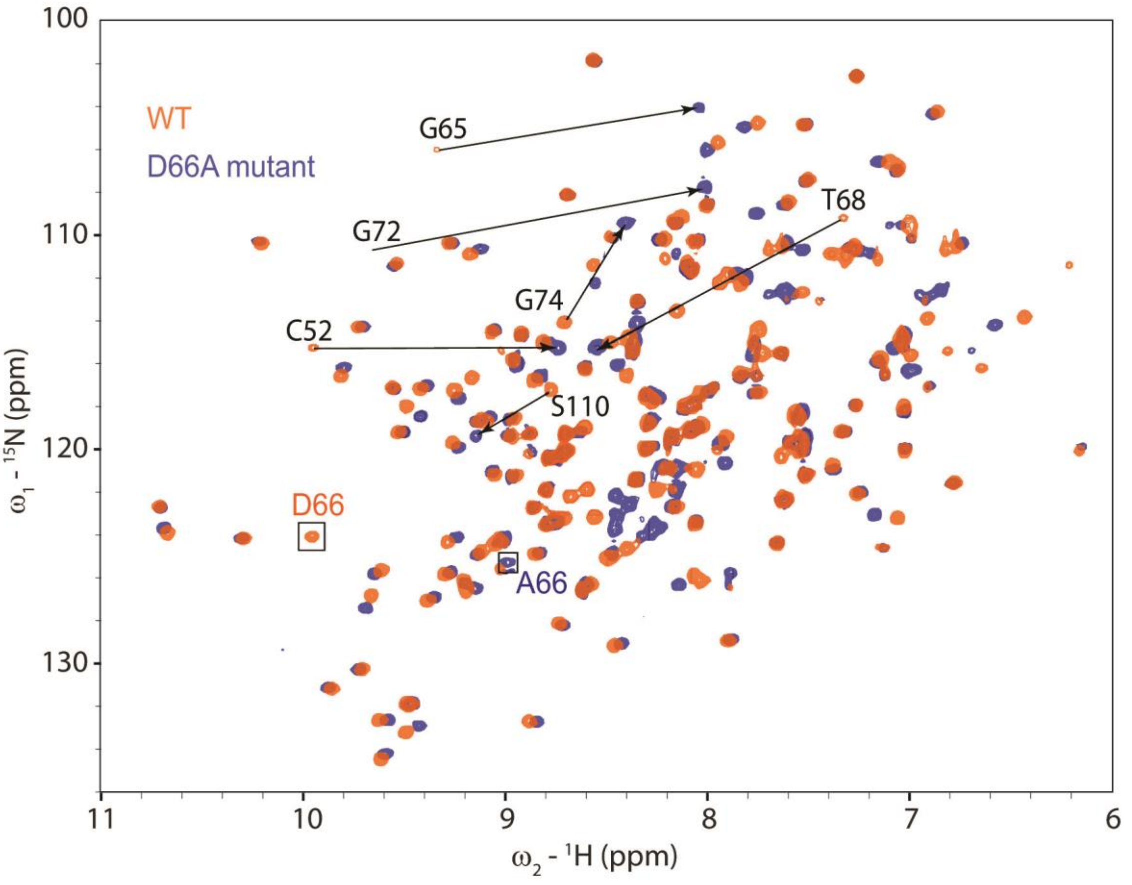


**Figure S8.** Comparison of CypA WT and D66A $^{15}N$-$^{1}H$ HSQC spectra with several residues showing large chemical shift perturbations (CSPs) highlighted. Most of these large CSPs occur in the 70s and 100s loops, showing a large difference in the ground state structure of the two proteins in this area.

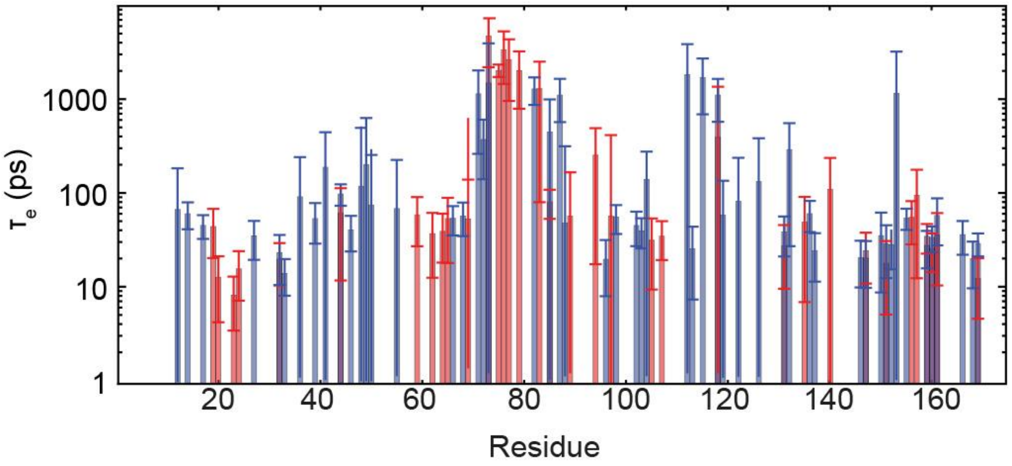


**Figure S9.** Using the relaxation data, a model-free analysis was carried out using the software package relax. $t_e$ is the effective correlation time for internal motion. Red denotes data for CypA and blue data for D66A. For residues in the 60 to 90 region the model free treatment fits $t_e$ values which approach the global tumbling time, and thus low values of generalized-order parameters derived from the model-free analysis are not indicative of fast motions.

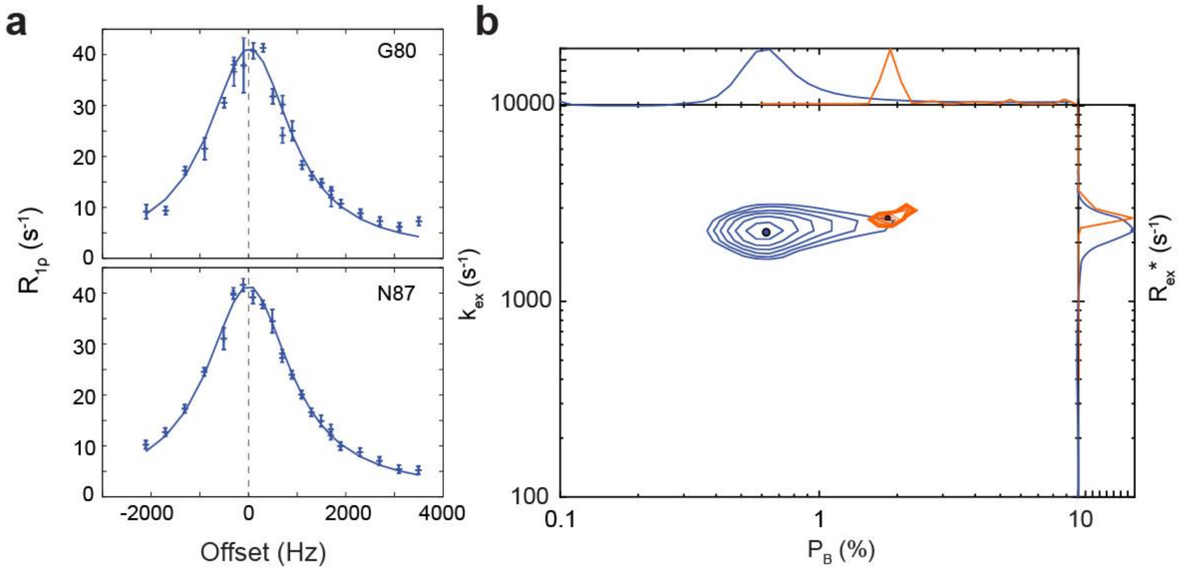


**Figure S10 a**. In addition to CPMG relaxation-dispersion, $R_{1\rho}$ experiments were carried out on the mutant protein as CPMG alone was insufficient to characterise exchange its parameters. By fitting the R1ρ data simultaneously with the CPMG dispersion profiles we could characterize the chemical exchange more precisely and also determine the signs for the chemical shift differences between the ground and excited state. **b.** Results of grid searching errors for fits of relaxation- dispersion data from the WT and D66A mutant. In both cases we find that we can fit the data well using a single $p_b$ and $k_{ex}$ for each protein. As seen in the results while the best fits of $k_{ex}$ are similar in both cases the population of the excited state in the mutant is significantly lower. It should be noted however that the 'excited' state appears to be different for the two proteins.

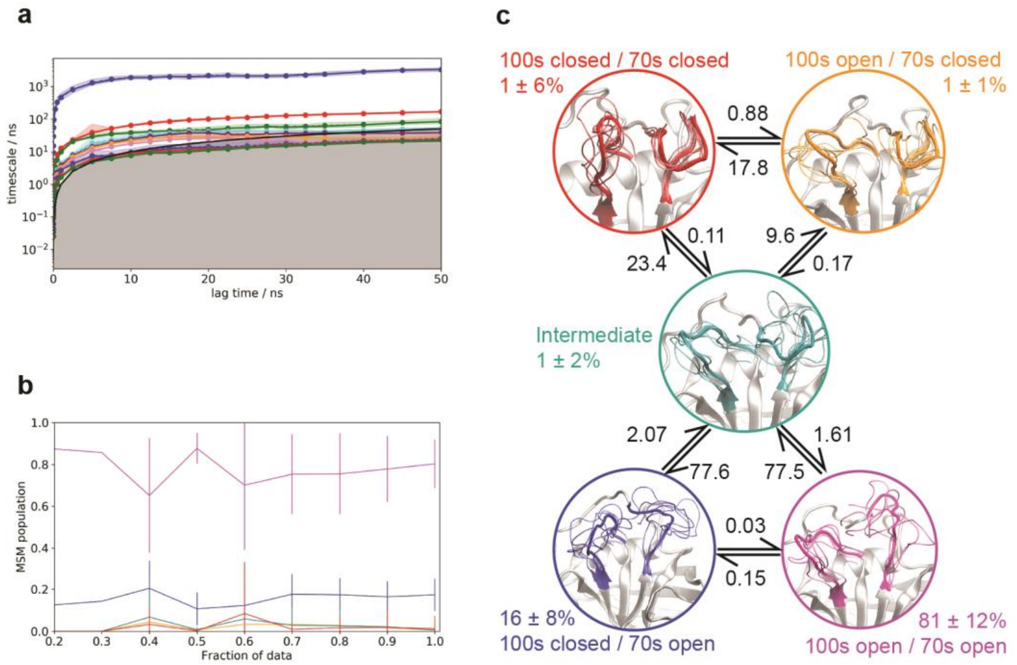


**Figure S11. a** Implied timescales of the MSM derived from the aMD trajectories as a function of lagtime. Shaded region represents the error as estimated by Bayesian analysis. **b** Predicted population of each MSM macrostate as a function of the fraction of the total dataset used to build the model. **c** Minimal set of sub-states that together contain the relevant amplitudes of motion and time-scales of state-to-state inter-conversion present in the full ensemble. The calculated rates and populations are indicated. Error bars on reported populations were obtained by bootstrapping of the MD trajectories assigned to the individual microstates.

### Supplementary Tables

**Table S1.** NMR assigment of CypA kindly provided by Dr. Xavier Hanoulle and Prof. Guy Lippens.

| | Chemical shift (ppm) | | | | | | | |
|---|---|---|---|---|---|---|---|---|
| **Residue** | **H$^{N}$** | **H$^{\alpha}$** | **H$^{\beta}$** | **other $^{1}$H** | **$^{15}$N** | **$^{13}$C$^{\alpha}$** | **$^{13}$C$^{\beta}$** | **other $^{13}$C, $^{15}$N** |
| **Met 1** | | 4.12 | 2.14 | H$^{\gamma}$ 2.58, 2.54; H$^{\varepsilon}$ 2.09 | | 54.8 | 32.9 | C$^{\gamma}$ 30.7; C$^{\varepsilon}$ 16.7 |
| **Val 2** | | 4.19 | 2.05 | H$^{\gamma}$ *0.94, 0.93* | | 61.5 | 33.1 | C$^{\gamma}$ *20.8, 20.2* |
| **Asn 3** | 8.64 | 5.14 | *2.77, 3.62* | H$^{\delta 2}$ 8.45, 6.98 | 126.5 | 51 | 37.7 | N$^{\delta 2}$ 109.7 |
| **Pro 4** | | 4.75 | *1.71, 2.36* | H$^{\gamma}$ *2.09, 2.19*; H$^{\delta}$ 4.09, 4.06 | | 62.7 | 33 | C$^{\gamma}$ 26.7; C$^{\delta}$ 50.6 |
| **Thr 5** | 8.76 | 5.72 | 4.05 | H$^{\gamma 2}$ 1.20 | 114.8 | 60.6 | 70.7 | C$^{\gamma}$ 22.1 |
| **Val 6** | 8.75 | 5.34 | 2.3 | H$^{\gamma}$ *1.05, 1.07* | 120.1 | 58.7 | 36.1 | C$^{\gamma}$ *21.7, 20.6* |
| **Phe 7** | 8.95 | 5.89 | 2.84, 2.63 | H$^{\delta}$ 6.73; H$^{\varepsilon}$ 6.99; H$^{\zeta}$ 7.06 | 119 | 55.7 | 43 | C$^{\delta}$ 132.0; C$^{\varepsilon}$ 130.5; C$^{\zeta}$ 129.5 |
| **Phe 8** | 9.53 | 5.32 | *3.46, 3.25* | H$^{\delta}$ 7.49; H$^{\varepsilon}$ 7.18; H$^{\zeta}$ 6.19 | 116.8 | 53.3 | 42.8 | C$^{\delta}$ 129.1; C$^{\varepsilon}$ 131.1; C$^{\zeta}$ 130.5 |
| **Asp 9** | 9.26 | 5.53 | 2.85, 2.52 | | 124 | 54.7 | 41.2 | |
| **Ile 10** | 9.02 | 5.17 | 2.06 | H$^{\gamma 1}$ 1.84, 1.66; H$^{\gamma 2}$ 0.96; H$^{\delta}$ 0.99 | 124.1 | 58.1 | 37.6 | C$^{\gamma 1}$ 27.0; C$^{\gamma 2}$ 18.4; C$^{\delta 1}$ 11.4 |
| **Ala 11** | 9.6 | 5.17 | 1.05 | | 132.2 | 50.7 | 22.6 | |
| **Val 12** | 8.93 | 4.51 | 1.78 | H$^{\gamma}$ *0.34, 0.65* | 118.2 | 60.3 | 33.1 | C$^{\gamma}$ *21.1, 21.6* |
| **Asp 13** | 9.83 | 4.31 | 3.09, 2.50 | | 130.7 | 55.5 | 39.8 | |
| **Gly 14** | 8.54 | 4.21, 3.42 | | | 101.5 | 45.1 | | |
| **Glu 15** | 8.03 | 4.84 | 2.09, 1.98 | H$^{\gamma}$ 2.29, 2.19 | 123 | 52.8 | 30.2 | C$^{\gamma}$ 35.5 |
| **Pro 16** | | 4.18 | *1.88, 2.32* | H$^{\gamma}$ 2.15, 1.98; H$^{\delta}$ 3.95, 3.72 | | 64.5 | 32.2 | C$^{\gamma}$ 27.5; C$^{\delta}$ 50.9 |
| **Leu 17** | 9.18 | 4.69 | 1.60, 1.41 | H$^{\gamma}$ 1.56; H$^{\delta}$ *0.98, 0.82* | 125.8 | 55.3 | 44.1 | C$^{\gamma}$ 27.4; C$^{\delta}$ *22.6, 25.7* |
| **Gly 18** | 7.24 | 4.26, 3.95 | | | 102.2 | 45 | | |
| **Arg 19** | 8.32 | 5.62 | 1.99, 1.62 | H$^{\gamma}$ 1.61, 1.47; H$^{\delta}$ 2.77, 2.67; H$^{\varepsilon}$ 9.01; H$^{\eta}$ 6.09 | 121 | 54.9 | 33.3 | C$^{\gamma}$ 26.9; C$^{\delta}$ 43.0; N$^{\varepsilon}$ 85.9; N$^{\eta}$ 70.4 |
| **Val 20** | 9.36 | 4.6 | 1.74 | H$^{\gamma}$ *0.51, 0.85* | 126.6 | 59.9 | 34.8 | C$^{\gamma}$ *22.3, 21.9* |
| **Ser 21** | 8.75 | 5.51 | 3.72, 3.56 | | 120.1 | 55.2 | 66.3 | |
| **Phe 22** | 9.51 | 5.24 | *2.30, 2.43* | H$^{\delta}$ 6.52; H$^{\varepsilon}$ 6.15; H$^{\zeta}$ 6.64 | 118.9 | 55.7 | 42.5 | C$^{\delta}$ 131.9; C$^{\varepsilon}$ 130.4; C$^{\zeta}$ 128.3 |
| **Glu 23** | 8.74 | 4.75 | 1.82, 1.68 | H$^{\gamma}$ 2.19, 1.80 | 122.9 | 54.9 | 31.6 | C$^{\gamma}$ 36.5 |
| **Leu 24** | 8.18 | 4.7 | *1.88, 1.23* | H$^{\gamma}$ 1.69; H$^{\delta}$ *1.10, 0.46* | 122.3 | 51.6 | 43.1 | C$^{\gamma}$ 26.9; C$^{\delta}$ *26.7, 23.0* |
| **Phe 25** | 8.81 | 5.11 | *2.67,* | H$^{\delta}$ 6.87; H$^{\varepsilon}$ 7.00; | 124.4 | 54.3 | 35.9 | C$^{\delta}$ 131.0; C$^{\varepsilon}$ 130.5; |

| | | | | | | | | |
|---|---|---|---|---|---|---|---|---|
| | | | *3.26* | H$^{\zeta}$ 6.95 | | | | C$^{\zeta}$ 130.6 |
| **Ala 26** | 8.42 | 3.77 | 1.5 | | 128.7 | 54.2 | 18.6 | |
| **Asp 27** | 9.05 | 4.24 | 2.84, 2.72 | | 114.1 | 55.2 | 38.9 | |
| **Lys 28** | 7.53 | 4.56 | 1.77 | H$^{\gamma}$ 1.38; H$^{\delta}$ 1.56, 1.41; H$^{\varepsilon}$ 2.82 | 117.8 | 56.2 | 35.9 | C$^{\gamma}$ 24.8; C$^{\delta}$ 28.4; C$^{\varepsilon}$ 41.7 |
| **Val 29** | 8.35 | 4.45 | 2.57 | H$^{\gamma}$ *0.99*, *1.30* | 114.2 | 58 | 31.7 | C$^{\gamma}$ *25.2, 20.6* |
| **Pro 30** | | 4.39 | *1.96, 2.61* | H$^{\gamma}$ 2.06, 2.10; H$^{\delta}$ 3.54, 3.09 | | 66.1 | 31.3 | C$^{\gamma}$ 27.2; C$^{\delta}$ 50.4 |
| **Lys 31** | 10.64 | 4 | 1.69, 1.54 | H$^{\gamma}$ 1.11, 0.89; H$^{\delta}$ 1.41, 1.31; H$^{\varepsilon}$ 2.06, 1.85 | 123.5 | 60.2 | 32.3 | C$^{\gamma}$ 26.4; C$^{\delta}$ 29.5; C$^{\varepsilon}$ 40.8 |
| **Thr 32** | 10.25 | 3.98 | 4.16 | H$^{\gamma 2}$ 0.92; H$^{\gamma 1}$ 7.76 | 123.7 | 67.3 | 68.4 | C$^{\gamma}$ 22.2 |
| **Ala 33** | 9.27 | 4.03 | 1.43 | | 125.4 | 55.8 | 18.2 | |
| **Glu 34** | 8.02 | 4.51 | *2.26, 1.75* | H$^{\gamma}$ 2.45, 2.03 | 117 | 57.8 | 28 | C$^{\gamma}$ 33.5 |
| **Asn 35** | 7.12 | 4.05 | *2.40, 2.93* | H$^{\delta 2}$ 8.98, 6.80 | 115.4 | 56.6 | 39.4 | N$^{\delta 2}$ 115.0 |
| **Phe 36** | 7 | 4.11 | *3.05, 2.95* | H$^{\delta}$ 6.42; H$^{\varepsilon}$ 5.56 | 117.6 | 61.4 | 40 | C$^{\varepsilon}$ 130.7 |
| **Arg 37** | 8.93 | 3.63 | 1.93, 1.83 | H$^{\gamma}$ 1.50, 1.01; H$^{\delta}$ 3.53, 2.81; H$^{\varepsilon}$ 8.89 | 120.8 | 60.3 | 29.9 | C$^{\gamma}$ 45.2; C$^{\delta}$ 41.0; N$^{\varepsilon}$ 79.0 |
| **Ala 38** | 8.68 | 4.07 | 1.24 | | 118.9 | 54.3 | 18.2 | |
| **Leu 39** | 8.15 | 3.74 | *1.36, −0.33* | H$^{\gamma}$ 1.75; H$^{\delta}$ *0.87, 0.53* | 120.5 | 57.1 | 40.9 | C$^{\gamma}$ 26.2; C$^{\delta}$ *26.7, 24.2* |
| **Ser 40** | 7.88 | 4.44 | 4.22, 3.52 | H$^{\gamma}$ 3.43 | 119 | 62.3 | 62.8 | |
| **Thr 41** | 7.97 | 4.28 | 4.47 | H$^{\gamma 2}$ 1.32; H$^{\gamma 1}$ 6.68 | 108.2 | 62.7 | 69.2 | C$^{\gamma}$ 22.4 |
| **Gly 42** | 7.57 | 3.86, 3.48 | | | 108.1 | 45.6 | | |
| **Glu 43** | 8 | 4.11 | *2.20, 2.04* | H$^{\gamma}$ 2.13, 1.79 | 118.5 | 58.4 | 29.9 | C$^{\gamma}$ 35.2 |
| **Lys 44** | 9.08 | 4.27 | 1.61, 0.89 | H$^{\gamma}$ 1.03; H$^{\delta}$ 1.44; H$^{\varepsilon}$ 2.82, 2.67 | 118.1 | 54.1 | 30.3 | C$^{\gamma}$ 24.8; C$^{\delta}$ 27.3; C$^{\varepsilon}$ 41.9 |
| **Gly 45** | 7.92 | 4.29, 3.54 | | | 105.3 | 44.4 | | |
| **Phe 46** | 6.41 | 4.62 | 3.11, 2.71 | H$^{\delta}$ 6.61; H$^{\varepsilon}$ 6.98; H$^{\zeta}$ 7.06 | 113.5 | 53.9 | 39.4 | C$^{\delta}$ 132.7; C$^{\varepsilon}$ 130.6; C$^{\zeta}$ 129.2 |
| **Gly 47** | 7.72 | 4.34, 2.52 | | | 104.4 | 45.3 | | |
| **Tyr 48** | 6.9 | 4.21 | 2.82, 2.64 | H$^{\delta}$ 6.75; H$^{\varepsilon}$ 6.13 | 113.5 | 57.1 | 38.4 | |
| **Lys 49** | 8.46 | 3.59 | 1.98, 1.59 | H$^{\gamma}$ 1.31, 1.24; H$^{\delta}$ 1.93, 1.76; H$^{\varepsilon}$ 3.23, 3.05 | 124.6 | 60.9 | 31.2 | C$^{\gamma}$ 25.3; C$^{\delta}$ 29.7; C$^{\varepsilon}$ 41.8 |
| **Gly 50** | 9.47 | 4.39, 3.69 | | | 117.6 | 45 | | |
| **Ser 51** | 8.38 | 4.66 | 4.31, 4.19 | H$^{\gamma}$ 4.89 | 116.2 | 58.7 | 64.6 | |
| **Cys 52** | 9.86 | 5.89 | 3.03 | | 114.8 | 56 | 31.7 | |
| **Phe 53** | 8.68 | 4.85 | *2.89, 2.63* | H$^{\delta}$ 6.89; H$^{\varepsilon}$ 7.18; | 122.8 | 58.1 | 38.7 | C$^{\delta}$ 131.4; C$^{\varepsilon}$ 130.8; |

| | | | | | | | | |
|---|---|---|---|---|---|---|---|---|
| | | | | H$^{\zeta}$ 7.36 | | | | C$^{\zeta}$ 128.2 |
| **His 54** | 7.62 | 4.71 | 3.27, 2.81 | H$^{\delta2}$ 6.84; H$^{\varepsilon1}$ 7.35 | 119.9 | 57 | 31.3 | C$^{\delta2}$ 118.2; C$^{\varepsilon1}$ 137.5 |
| **Arg 55** | 7.04 | 5.1 | 1.38, 1.25 | H$^{\gamma}$ 1.56, 1.21; H$^{\delta}$ 3.16, 3.12; H$^{\varepsilon}$ 7.11 | 122.8 | 54.7 | 33.7 | C$^{\gamma}$ 28.0; C$^{\delta}$ 43.4; N$^{\varepsilon}$ 83.9 |
| **Ile 56** | 9.15 | 4.59 | 1.57 | H$^{\gamma1}$ 1.55, 0.97; H$^{\gamma2}$ 1.00; H$^{\delta}$ 0.85 | 126.2 | 61.5 | 41.6 | C$^{\gamma1}$ 27.1; C$^{\gamma2}$ 17.6; C$^{\delta1}$ 14.4 |
| **Ile 57** | 8.71 | 5.19 | 1.9 | H$^{\gamma1}$ 1.45, 1.14; H$^{\gamma2}$ 1.11; H$^{\delta}$ 0.91 | 127.7 | 57.5 | 40 | C$^{\gamma1}$ 27.2; C$^{\gamma2}$ 17.5; C$^{\delta1}$ 13.4 |
| **Pro 58** | | 4.32 | *1.80, 2.19* | H$^{\gamma}$ 2.03, 1.86; H$^{\delta}$ 4.00, 3.44 | | 62.9 | 31.9 | C$^{\gamma}$ 26.9; C$^{\delta}$ 52.1 |
| **Gly 59** | 9.68 | 4.03, 3.79 | | | 113.9 | 44.9 | | |
| **Phe 60** | 8.15 | 5.08 | 3.15, 3.03 | H$^{\delta}$ 7.26; H$^{\varepsilon}$ 7.14 | 119 | 56.7 | 39.6 | C$^{\delta}$ 130.3; C$^{\varepsilon}$ 131.2 |
| **Met 61** | 8.08 | 5.31 | *1.39, 2.14* | H$^{\gamma}$ 1.00; H$^{\varepsilon}$ 1.25 | 111 | 54.8 | 34.7 | C$^{\gamma}$ 28.9; C$^{\varepsilon}$ 18.3 |
| **Cys 62** | 8.45 | 4.87 | 2.96, 2.48 | H$^{\gamma}$ 1.60 | 114.6 | 57.3 | 30 | |
| **Gln 63** | 9.64 | 5.38 | 2.15, 1.79 | H$^{\gamma}$ 2.53, 1.98; H$^{\varepsilon2}$ 6.96, 6.96 | 126.5 | 54.4 | 30.4 | C$^{\gamma}$ 33.1; N$^{\varepsilon2}$ 109.1 |
| **Gly 64** | 7.36 | 4.71, 2.57 | | | 110.5 | 44.4 | | |
| **Gly 65** | 9.29 | *3.78, 4.77* | | | 105.6 | 46.4 | | |
| **Asp 66** | 9.91 | 4.29 | 2.93, 1.70 | | 123.7 | 51.5 | 38.7 | |
| **Phe 67** | 6.62 | 4.6 | 3.92, 2.57 | H$^{\delta}$ 7.18; H$^{\varepsilon}$ 7.32; H$^{\zeta}$ 7.41 | 115.9 | 55.8 | 39.3 | C$^{\delta}$ 132.3; C$^{\varepsilon}$ 131.7; C$^{\zeta}$ 128.5 |
| **Thr 68** | 7.32 | 4.64 | 4.09 | H$^{\gamma2}$ 0.68 | 108.9 | 61.6 | 68.7 | C$^{\gamma}$ 22.8 |
| **Arg 69** | 8.65 | 4.35 | *1.87, 1.60* | H$^{\gamma}$ 1.46, 1.37; H$^{\delta}$ 3.05, 2.94; H$^{\varepsilon}$ 7.01 | 121.8 | 55 | 31.6 | C$^{\gamma}$ 26.5; C$^{\delta}$ 42.7; N$^{\varepsilon}$ 84.4 |
| **His 70**[b] | 6.64 | 4.39 | 3.22 | H$^{\delta2}$ 6.65; H$^{\varepsilon1}$ 7.68 | 111 | 57.1 | 28.4 | C$^{\delta2}$ 118.0; C$^{\varepsilon1}$ 138.1 |
| **Asn 71** | 7.52 | 4.59 | 3.14, 2.65 | H$^{\delta2}$ 7.63, 6.78 | 112.3 | 52.2 | 38.7 | N$^{\delta2}$ 110.3 |
| **Gly 72** | 9.62 | 4.40, 3.16 | | | 110.2 | 44.6 | | |
| **Thr 73** | 7.91 | 4.48 | 4.25 | H$^{\gamma2}$ 1.02; H$^{\gamma1}$ 5.77 | 111.8 | 62.1 | 70.3 | C$^{\gamma}$ 20.8 |
| **Gly 74** | 8.67 | 4.46, 3.46 | | | 113.7 | 44.8 | | |
| **Gly 75** | 8.07 | 4.55, 2.54 | | | 108.8 | 43 | | |
| **Lys 76** | 6.98 | 4.59 | *1.54, 1.88* | H$^{\gamma}$ 0.98, 0.91; H$^{\delta}$ 1.19; H$^{\varepsilon}$ 2.17, 1.53 | 115.5 | 55.7 | 34.5 | C$^{\gamma}$ 23.3; C$^{\delta}$ 29.2; C$^{\varepsilon}$ 41.0 |
| **Ser 77** | 7.76 | 5.18 | 4.25 | H$^{\gamma}$ 5.10 | 114.1 | 56.8 | 69.1 | |
| **Ile 78** | 8.53 | 4.17 | 1.7 | H$^{\gamma1}$ 0.64, –0.47; H$^{\gamma2}$ 0.71; H$^{\delta}$ 0.47 | 111 | 63.5 | 37.3 | C$^{\gamma1}$ 23.0; C$^{\gamma2}$ 17.5; C$^{\delta1}$ 14.1 |
| **Tyr 79** | 8.01 | 4.67 | *2.33, 3.42* | H$^{\delta}$ 6.41; H$^{\varepsilon}$ 6.08 | 120.5 | 56.1 | 38.6 | C$^{\delta}$ 132.5; C$^{\varepsilon}$ 117.7 |
| **Gly 80** | 7.11 | 4.60, | | | 106.4 | 43.8 | | |

| | | | | | | | | |
|---|---|---|---|---|---|---|---|---|
| | | 3.79 | | | | | | |
| **Glu 81** | 8.88 | 3.96 | 2.07 | H$^{\gamma}$ 2.34, 2.23 | 122.8 | 60.1 | 30.1 | C$^{\gamma}$ 36.3 |
| **Lys 82** | 7.84 | 5.51 | 1.63, 1.56 | H$^{\gamma}$ 1.25; H$^{\delta}$ 1.51, 1.55; H$^{\varepsilon}$ 2.90 | 112.6 | 53.7 | 36.3 | C$^{\gamma}$ 24.3; C$^{\delta}$ 29.0; C$^{\varepsilon}$ 42.0 |
| **Phe 83** | 9.16 | 4.95 | 3.02 | H$^{\delta}$ 7.33; H$^{\varepsilon}$ 7.51; H$^{\zeta}$ 7.38 | 116.4 | 55.7 | 41.3 | C$^{\delta}$ 134.2; C$^{\varepsilon}$ 130.8; C$^{\zeta}$ 128.5 |
| **Glu 84** | 9.25 | 3.8 | 2.02, 1.82 | H$^{\gamma}$ 2.28, 2.04 | 119.4 | 56.2 | 29.1 | C$^{\gamma}$ 35.2 |
| **Asp 85** | 8.57 | 4.23 | 2.40, 1.91 | | 118.6 | 54.1 | 40.8 | |
| **Glu 86** | 9.44 | 3.79 | 2.12, 1.83 | H$^{\gamma}$ 2.36 | 131.5 | 60.2 | 31.6 | C$^{\gamma}$ 35.1 |
| **Asn 87** | 7.03 | 4.13 | *2.73, 3.44* | H$^{\delta2}$ 7.93, 6.88 | 106.5 | 52.6 | 39.3 | N$^{\delta2}$ 116.7 |
| **Phe 88** | 8.31 | 5.93 | 3.52, 2.53 | H$^{\delta}$ 7.25; H$^{\varepsilon}$ 6.86; H$^{\zeta}$ 6.85 | 112.7 | 55.7 | 38.7 | C$^{\delta}$ 131.9; C$^{\varepsilon}$ 130.6; C$^{\zeta}$ 128.7 |
| **Ile 89** | 8.27 | 3.65 | 1.54 | H$^{\gamma1}$ 1.48, 1.14; H$^{\gamma2}$ 0.84; H$^{\delta}$ 0.90 | 119.6 | 64.8 | 38.9 | C$^{\gamma1}$ 29.2; C$^{\gamma2}$ 16.8; C$^{\delta1}$ 13.6 |
| **Leu 90** | 7.73 | 4.45 | *1.82, 1.36* | H$^{\gamma}$ 1.45; H$^{\delta}$ *0.75, 0.64* | 117 | 53.9 | 41.8 | C$^{\gamma}$ 27.0; C$^{\delta}$ *26.8, 22.5* |
| **Lys 91** | 8.05 | 4.63 | *1.54, 2.26* | H$^{\gamma}$ 1.44, 1.34; H$^{\delta}$ 1.62, 1.56; H$^{\varepsilon}$ 3.00 | 118.7 | 54.3 | 34.5 | C$^{\gamma}$ 25.7; C$^{\delta}$ 28.1; C$^{\varepsilon}$ 42.3 |
| **His 92** | 10.67 | 4.34 | *2.88, 3.14* | H$^{\delta1}$ 12.21; H$^{\delta2}$ 6.74; H$^{\varepsilon1}$ 8.05 | 122.2 | 56.8 | 25.9 | C$^{\delta2}$ 118.2; C$^{\varepsilon1}$ 139.7; N$^{\delta1}$ 169.1 |
| **Thr 93** | 7.25 | 4 | 4.43 | H$^{\gamma2}$ 1.20 | 110.2 | 63 | 69.3 | C$^{\gamma}$ 22.5 |
| **Gly 94** | 7.47 | 4.36, 3.64 | | | 107 | 45.4 | | |
| **Pro 95** | | 3.81 | *1.84, 2.01* | H$^{\gamma}$ 2.12, 1.83; H$^{\delta}$ 3.60, 3.39 | | 62.5 | 32 | C$^{\gamma}$ 27.3; C$^{\delta}$ 49.6 |
| **Gly 96** | 9.25 | 4.56, 3.33 | | | 109.9 | 44.8 | | |
| **Ile 97** | 6.76 | 4.11 | 2.22 | H$^{\gamma1}$ 1.80, 1.64; H$^{\gamma2}$ 0.98; H$^{\delta}$ 0.96 | 121.2 | 59.3 | 37.8 | C$^{\gamma1}$ 28.9; C$^{\gamma2}$ 20.2; C$^{\delta1}$ 9.0 |
| **Leu 98** | 7.87 | 5 | *0.15, 0.08* | H$^{\gamma}$ 0.41; H$^{\delta}$ *−0.17, −0.59* | 128.5 | 53.2 | 43.5 | C$^{\gamma}$ 26.4; C$^{\delta}$ *24.1, 25.1* |
| **Ser 99** | 8.29 | 5.32 | 3.23, 2.88 | | 118.3 | 54.9 | 65.3 | |
| **Met 100** | 8.55 | 5.26 | 2.57, 2.29 | H$^{\gamma}$ 2.79, 2.43; H$^{\varepsilon}$ 2.08 | 122.8 | 53.7 | 31 | C$^{\gamma}$ 32.0; C$^{\varepsilon}$ 16.1 |
| **Ala 101** | 8.03 | 4.26 | 1.1 | | 126 | 51.4 | 19.4 | |
| **Asn 102** | 8.14 | 4.57 | 3.24, 2.89 | H$^{\delta2}$ 7.89, 6.16 | 113.2 | 53.8 | 40.4 | N$^{\delta2}$ 119.4 |
| **Ala 103** | 8.77 | 4.78 | 1.3 | | 123.1 | 50.2 | 19 | |
| **Gly 104** | 8.11 | 4.59, 3.70 | | | 108.9 | 43.3 | | |
| **Pro 105** | | 4.32 | *1.80, 2.32* | H$^{\gamma}$ 2.11, 1.95; H$^{\delta}$ 3.60, 3.51 | | 63.8 | 31.9 | C$^{\gamma}$ 27.6; C$^{\delta}$ 49.2 |
| **Asn 106** | 8.86 | 3.99 | *3.08, 2.66* | H$^{\delta2}$ 7.72, 7.08 | 118.9 | 54 | 36.9 | N$^{\delta2}$ 116.0 |
| **Thr 107** | 10.16 | 4.43 | 4.38 | H$^{\gamma2}$ 0.88; H$^{\gamma1}$ 4.68 | 110 | 60 | 68.8 | C$^{\gamma}$ 21.0 |
| **Asn 108** | 7.37 | 4.21 | 1.54, 0.92 | H$^{\delta2}$ 7.41, 5.76 | 120.3 | 55.4 | 39.6 | N$^{\delta2}$ 112.7 |

| | | | | | | | | |
|---|---|---|---|---|---|---|---|---|
| **Gly 109** | 9.14 | 4.59, 3.62 | | | 110.6 | 45.5 | | |
| **Ser 110** | 8.74 | 4.7 | 4.60, 4.37 | $H^{\gamma}$ 4.93 | 116.8 | 57.5 | 66 | |
| **Gln 111** | 8.39 | 5.17 | 2.20, 2.07 | $H^{\gamma}$ 2.44, 1.84; $H^{\varepsilon 2}$ 7.30, 6.19 | 124.3 | 57.5 | 31.6 | $C^{\gamma}$ 35.2; $N^{\varepsilon 2}$ 111.0 |
| **Phe 112** | 8.06 | 5.7 | 3.28 | $H^{\delta}$ 6.90, 7.80; $H^{\varepsilon}$ 6.19, 7.22; $H^{\zeta}$ 6.75 | 117.6 | 55.4 | 43 | $C^{\delta}$ 133.9 |
| **Phe 113** | 9.79 | 5.66 | 2.89, 2.83 | $H^{\delta}$ 6.82; $H^{\varepsilon}$ 6.75; $H^{\zeta}$ 6.88 | 116.2 | 54.3 | 42.6 | $C^{\delta}$ 131.6; $C^{\varepsilon}$ 130.4; $C^{\zeta}$ 129.3 |
| **Ile 114** | 9.1 | 4.79 | 1.8 | $H^{\gamma 1}$ 2.09, 1.09; $H^{\gamma 2}$ 0.83; $H^{\delta}$ 1.20 | 118.4 | 60.1 | 40.1 | $C^{\gamma 1}$ 27.2; $C^{\gamma 2}$ 17.7; $C^{\delta 1}$ 15.0 |
| **Cys 115** | 9.59 | 4.75 | *3.09, 3.45* | | 125.3 | 60.7 | 28.5 | |
| **Thr 116** | 8.93 | 4.32 | 4.42 | $H^{\gamma 2}$ 0.82 | 115.5 | 60.6 | 66.5 | $C^{\gamma}$ 23.0 |
| **Ala 117** | 7.6 | 4.36 | 1.41 | | 122 | 50.4 | 22.6 | |
| **Lys 118** | 8.66 | 3.78 | 1.96, 1.63 | $H^{\gamma}$ 1.50; $H^{\delta}$ 1.74; $H^{\varepsilon}$ 3.01 | 119.6 | 57.7 | 32.5 | $C^{\gamma}$ 24.6; $C^{\delta}$ 29.5; $C^{\varepsilon}$ 41.7 |
| **Thr 119** | 7.31 | 3.62 | 2.81 | $H^{\gamma 2}$ 0.97; $H^{\gamma 1}$ 3.07 | 118.8 | 57 | 68.5 | $C^{\gamma}$ 19.7 |
| **Glu 120** | 9.09 | 4.11 | *2.18, 2.11* | $H^{\gamma}$ 2.50, 2.39 | 124.4 | 59.1 | 29 | $C^{\gamma}$ 35.3 |
| **Trp 121** | 7.24 | 4.65 | *3.37, 3.32* | $H^{\delta 1}$ 7.02; $H^{\varepsilon 1}$ 9.65; $H^{\varepsilon 3}$ 7.66; $H^{\zeta 3}$ 7.32; $H^{\eta 2}$ 7.11; $H^{\zeta 2}$ 6.61 | 117.5 | 59.5 | 26.8 | $C^{\delta 1}$ 127.7; $C^{\varepsilon 3}$ 119.0; $C^{\zeta 2}$ 114.9; $C^{\zeta 3}$ 123.7; $C^{\eta 2}$ 125.1; $N^{\varepsilon 1}$ 129.8 |
| **Leu 122** | 6.99 | 4.24 | *0.10, 1.13* | $H^{\gamma}$ 0.37; $H^{\delta}$ *0.37, 0.65* | 119.5 | 54.5 | 38.9 | $C^{\gamma}$ 26.0; $C^{\delta}$ *25.2, 21.5* |
| **Asp 123** | 7.6 | 5.16 | 2.93, 2.68 | | 121.8 | 55.6 | 39.3 | |
| **Gly 124** | 9.49 | 3.89, 2.88 | | | 110.8 | 44.8 | | |
| **Lys 125** | 7.68 | 4.11 | 1.82, 1.67 | $H^{\gamma}$ 1.25, 1.22; $H^{\delta}$ 1.67; $H^{\varepsilon}$ 3.00 | 115 | 56.1 | 35.4 | $C^{\gamma}$ 25.1; $C^{\delta}$ 29.0; $C^{\varepsilon}$ 42.0 |
| **His 126** | 7.58 | 4.8 | *3.18, 3.99* | $H^{\delta 2}$ 6.95; $H^{\varepsilon 1}$ 8.22 | 119.2 | 54.6 | 31.5 | $C^{\delta 2}$ 118.1 |
| **Val 127** | 8.47 | 4.26 | 2 | $H^{\gamma}$ *1.13, 1.02* | 124.7 | 63.6 | 33.4 | $C^{\gamma}$ *20.9, 21.1* |
| **Val 128** | 9.48 | 4.16 | 1.71 | $H^{\gamma}$ *0.93, 0.08* | 132.8 | 62.9 | 31.4 | $C^{\gamma}$ *21.7, 19.9* |
| **Phe 129** | 8.07 | 5.25 | 3.10, 2.49 | $H^{\delta}$ 6.52, 7.48; $H^{\varepsilon}$ 5.64, 6.81; $H^{\zeta}$ 6.43 | 117.5 | 55.9 | 42.4 | $C^{\delta}$ 130.6; $C^{\varepsilon}$ 131.2 |
| **Gly 130** | 7.29 | 3.16, 2.98 | | | 110.5 | 46.3 | | |
| **Lys 131** | 8.35 | 5.2 | 1.82 | $H^{\gamma}$ 1.50, 1.04; $H^{\delta}$ 1.76, 1.68; $H^{\varepsilon}$ 2.96 | 115.1 | 54.7 | 35.9 | $C^{\gamma}$ 24.0; $C^{\delta}$ 29.3; $C^{\varepsilon}$ 42.0 |
| **Val 132** | 9.01 | 3.89 | 1.91 | $H^{\gamma}$ *0.59, 1.03* | 123.8 | 63.9 | 32.5 | $C^{\gamma}$ *21.2, 22.9* |
| **Lys 133** | 9.44 | 4.42 | 1.71, 1.63 | $H^{\gamma}$ 1.44, 1.30; $H^{\delta}$ 1.62; $H^{\varepsilon}$ 2.95 | 131.3 | 56.7 | 33.8 | $C^{\gamma}$ 24.6; $C^{\delta}$ 29.4; $C^{\varepsilon}$ 41.3 |
| **Glu 134** | 7.53 | 4.56 | 1.95, 1.85 | $H^{\gamma}$ 2.18, 2.14 | 118.2 | 55.1 | 32.7 | $C^{\gamma}$ 36.6 |
| **Gly 135** | 8.65 | 4.73, 3.98 | | | 107.8 | 45.8 | | |

| | | | | | | | | |
|---|---|---|---|---|---|---|---|---|
| **Met 136** | 8.82 | 4.44 | *1.92, 2.09* | H$^{\gamma}$ 2.75, 2.59; H$^{\varepsilon}$ 1.98 | 122.3 | 56.7 | 30 | C$^{\gamma}$ 32.7; C$^{\varepsilon}$ 17.1 |
| **Asn 137** | 8.89 | 4.45 | 2.82, 2.76 | H$^{\delta2}$ 7.73, 7.00 | 114.2 | 55.9 | 35.8 | N$^{\delta2}$ 114.6 |
| **Ile 138** | 7.63 | 3.75 | 2.44 | H$^{\gamma1}$ 1.51, 1.27; H$^{\gamma2}$ 0.63; H$^{\delta}$ 0.65 | 123.9 | 61.3 | 34.3 | C$^{\gamma1}$ 27.0; C$^{\gamma2}$ 17.2; C$^{\delta1}$ 9.1 |
| **Val 139** | 7.24 | 3.84 | 2.44 | H$^{\gamma}$ *0.80, 1.02* | 121.7 | 66 | 31 | C$^{\gamma}$ *22.1, 23.1* |
| **Glu 140** | 8.27 | 4.04 | 2.06, 1.96 | H$^{\gamma}$ 2.40, 2.20 | 117.1 | 59.1 | 29.4 | C$^{\gamma}$ 36.8 |
| **Ala 141** | 7.49 | 4.06 | 1.58 | | 120.8 | 54.7 | 18 | |
| **Met 142** | 8.25 | 3.93 | *2.51, 1.93* | H$^{\gamma}$ *2.26, 2.62*; H$^{\varepsilon}$ 1.16 | 117.3 | 59.5 | 33.6 | C$^{\gamma}$ 31.1; C$^{\varepsilon}$ 14.4 |
| **Glu 143** | 7.81 | 3.71 | 2.14, 1.95 | H$^{\gamma}$ *2.04, 2.34* | 116.1 | 59.1 | 30.3 | C$^{\gamma}$ 37.7 |
| **Arg 144** | 7.02 | 3.91 | 1.45, 1.36 | H$^{\gamma}$ 1.58, 1.46; H$^{\delta}$ 3.04; H$^{\varepsilon}$ 7.10 | 114.4 | 57.4 | 29.3 | C$^{\gamma}$ 27.5; C$^{\delta}$ 43.1; N$^{\varepsilon}$ 84.8 |
| **Phe 145** | 7.6 | 4.79 | *2.90, 3.53* | H$^{\delta}$ 7.30; H$^{\varepsilon}$ 7.16; H$^{\zeta}$ 7.27 | 115.1 | 57.6 | 38.9 | C$^{\delta}$ 131.6; C$^{\varepsilon}$ 129.2; C$^{\zeta}$ 129.8 |
| **Gly 146** | 7.5 | *4.61, 3.62* | | | 104.5 | 43.9 | | |
| **Ser 147** | 8.18 | 4.75 | 4.15, 4.04 | | 109.8 | 58 | 65.5 | |
| **Arg 148** | 8.8 | 4.16 | 1.92, 1.88 | H$^{\gamma}$ 1.74, 1.69; H$^{\delta}$ 3.22; H$^{\varepsilon}$ 7.24 | 119.8 | 59.5 | 29.7 | C$^{\gamma}$ 26.9; C$^{\delta}$ 43.0; N$^{\varepsilon}$ 83.8 |
| **Asn 149** | 7.84 | 4.84 | 3.16, 2.80 | H$^{\delta2}$ 7.53, 6.72 | 111.4 | 52.1 | 38.1 | N$^{\delta2}$ 110.0 |
| **Gly 150** | 8.03 | 4.10, 3.93 | | | 109.9 | 44.8 | | |
| **Lys 151** | 7.5 | 4.41 | 1.87, 1.82 | H$^{\gamma}$ 1.53, 1.48; H$^{\delta}$ 1.77, 1.74; H$^{\varepsilon}$ 3.05 | 119.6 | 57 | 32.3 | C$^{\gamma}$ 24.8; C$^{\delta}$ 28.8; C$^{\varepsilon}$ 41.9 |
| **Thr 152** | 8.81 | 5.53 | 4.63 | H$^{\gamma2}$ 1.34; H$^{\gamma1}$ 5.01 | 116.3 | 59.8 | 71.1 | C$^{\gamma}$ 22.1 |
| **Ser 153** | 9.39 | 4.45 | 4.14, 4.02 | | 116.8 | 58.7 | 63.5 | |
| **Lys 154** | 7.52 | 4.53 | 1.41, 0.83 | H$^{\gamma}$ 1.42, 1.18; H$^{\delta}$ 1.44, 1.13; H$^{\varepsilon}$ 2.98, 2.89 | 119.1 | 54.4 | 38.3 | C$^{\gamma}$ 26.2; C$^{\delta}$ 29.2; C$^{\varepsilon}$ 42.3 |
| **Lys 155** | 8.78 | 4.31 | *1.78, 1.87* | H$^{\gamma}$ 1.40, 1.31; H$^{\delta}$ 1.66; H$^{\varepsilon}$ 2.92, 2.90 | 121.5 | 56.1 | 32 | C$^{\gamma}$ 24.8; C$^{\delta}$ 28.8; C$^{\varepsilon}$ 41.5 |
| **Ile 156** | 9.57 | 5.09 | 2.34 | H$^{\gamma1}$ 1.48, 1.44; H$^{\gamma2}$ 0.92; H$^{\delta}$ 0.64 | 134 | 59.2 | 36.8 | C$^{\gamma1}$ 26.7; C$^{\gamma2}$ 19.0; C$^{\delta1}$ 13.4 |
| **Thr 157** | 9.23 | 5.26 | 3.88 | H$^{\gamma2}$ 0.99 | 117 | 59.4 | 72.5 | C$^{\gamma}$ 22.4 |
| **Ile 158** | 8.57 | 4.24 | 1.67 | H$^{\gamma1}$ 1.41, 0.34; H$^{\gamma2}$ 0.31; H$^{\delta}$ –0.14 | 121.5 | 60.8 | 36.8 | C$^{\gamma1}$ 28.3; C$^{\gamma2}$ 18.3; C$^{\delta1}$ 13.7 |
| **Ala 159** | 8.86 | 4.12 | 1.29 | | 132.3 | 54 | 19 | |
| **Asp 160** | 8.06 | 4.89 | 2.96, 2.86 | | 111.3 | 52.6 | 43.5 | |
| **Cys 161** | 8.59 | 4.54 | 3.22, 3.09 | | 115.8 | 55.1 | 31.1 | |
| **Gly 162** | 6.83 | 3.89, 3.55 | | | 103.9 | 45.3 | | |

| | | | | | | | | |
|---|---|---|---|---|---|---|---|---|
| **Gln 163** | 9.04 | 5.07 | 1.96, 1.75 | H$^{\gamma}$ 2.57, 2.47; H$^{\varepsilon 2}$ 8.14, 7.16 | 120.7 | 54.8 | 30.6 | C$^{\gamma}$ 33.2; N$^{\varepsilon 2}$ 110.6 |
| **Leu 164** | 8.59 | 4.61 | 1.62 | H$^{\gamma}$ 1.65; H$^{\delta}$ *0.83, 0.74* | 125.8 | 54.7 | 43.3 | C$^{\gamma}$ 27.1; C$^{\delta}$ *25.2, 23.2* |
| **Glu 165** | 8.1 | 4.16 | 2.09, 1.92 | H$^{\gamma}$ 2.20 | 125.9 | 57.9 | 31.6 | C$^{\gamma}$ 36.8 |

**Table S2.** X-ray refinement statistics

| | |
|---|---|
| **Protein** | CypA |
| **Ligand** | 1 |
| **PDB ID** | 6GS6 |
| **Data collection and processing** | |
| High resolution limit | 1.16 (1.16) |
| Low resolution limit | 54.57 (1.18) |
| Completeness | 99.64 (99.27) |
| Multiplicity | 9.1 (6.5) |
| I/sigma | 14.2 (1.0) |
| Rmerge | 0.048 (1.70) |
| Unit cell dimensions: | |
| a (Å) | 63.01 |
| b (Å) | 63.01 |
| c (Å) | 94.72 |
| α (°) | 90.00 |
| β (°) | 90.00 |
| γ (°) | 120.00 |
| Space group | P $3_2$ 2 1 |
| Total observations | 686777 (23855) |
| Total unique | 75550 (3671) |
| **Refinement statistics** | |
| No. of residues per chain (No. of chains) | 165 |
| No. of ligands | 1 |
| No. of waters | 83 |
| Percentage of free reflections | 5.20 |
| R factor | 0.15 |
| R free | 0.20 |
| Rms BondLength | 0.01 |
| Rms BondAngle | 1.43 |
| **Ramachandran plot** | |
| Favoured (%) | 96.7 |
| Allowed (%) | 3.3 |
| No. of outliers* | 0 |

**Table S3.** Exchange parameters for WT and D66A obtained from a two state exchange model.

| | WT | |
|---|---|---|
| **Residue** | **Δω** | **Err** |
| **Glu 15** | 0.2 | 0.0 |
| **Leu 24** | 0.6 | 0.1 |
| **Thr 32** | 2.2 | 0.2 |
| **Phe 46** | 1.0 | 0.2 |
| **Tyr 48** | 1.1 | 0.2 |
| **Asp 66** | 3.0 | 0.2 |
| **Thr 68** | 6.3 | 0.1 |
| **His 70** | 1.1 | 0.2 |
| **Asn 71** | 5.9 | 0.2 |
| **Thr 73** | 1.3 | 0.2 |
| **Gly 74** | 5.9 | 0.2 |
| **Ser 77** | 3.9 | 0.2 |
| **Gly 80** | 1.0 | 0.2 |
| **Asn 87** | 1.1 | 0.2 |
| **Ile 89** | 1.4 | 0.2 |
| **Gly 96** | 1.4 | 0.2 |
| **Ser 99** | 1.8 | 0.2 |
| **Asn 102** | 1.9 | 0.2 |
| **Ser 110** | 3.0 | 0.2 |
| **Gln 111** | 1.8 | 0.2 |
| **Phe 113** | 1.4 | 0.2 |
| **Glu 120** | 1.4 | 0.2 |
| **Leu 122** | 1.4 | 0.2 |
| **Lys 131** | 1.4 | 0.2 |
| **$k_{ex}$($s^{-1}$)** | 2150 | 90 |
| **pb(frac)** | 0.021 | 0.001 |
| | **D66A** | |
| **Residue** | **Δω** | **Err** |
| **Ala 33** | 2.5 | 0.3 |
| **Arg 55** | 0.8 | 0.3 |
| **Gly 72** | 2.4 | 0.2 |
| **Gly 74** | -2.2 | 0.2 |
| **Gly 80** | -7.6 | 0.2 |
| **Phe 83** | 3.8 | 0.2 |
| **Asn 87** | 2.2 | 0.2 |
| **Glu 120** | 3.9 | 0.1 |
| **kex($s^{-1}$)** | 2200 | 350 |
| **pb(frac)** | 0.007 | 0.001 |

**Table S4.** Measured Rex values for WT, D66A. The 70s-loop region is highlighted.

| WT | | D66A | |
|---|---|---|---|
| Residue | ($s^{-1}$) | Residue | ($s^{-1}$) |
| **Met 1** | ND | **Met 1** | ND |
| **Val 2** | ND | **Val 2** | 0.0 |
| **Asn 3** | ND | **Asn 3** | ND |
| **Thr 5** | 0.2 | **Thr 5** | 0.4 |
| **Val 6** | ND | **Val 6** | ND |
| **Phe 7** | ND | **Phe 7** | ND |
| **Phe 8** | ND | **Phe 8** | 1.0 |
| **Asp 9** | 1.1 | **Asp 9** | 1.2 |
| **Ile 10** | 1.3 | **Ile 10** | 0.3 |
| **Ala 11** | 1.3 | **Ala 11** | 2.7 |
| **Val 12** | ND | **Val 12** | ND |
| **Asp 13** | 2.2 | **Asp 13** | ND |
| **Gly 14** | 2.7 | **Gly 14** | 0.7 |
| **Glu 15** | 1.6 | **Glu 15** | 1.0 |
| **Leu 17** | ND | **Leu 17** | 2.4 |
| **Gly 18** | 3.6 | **Gly 18** | 1.0 |
| **Arg 19** | ND | **Arg 19** | 1.3 |
| **Val 20** | ND | **Val 20** | 2.5 |
| **Ser 21** | 0.4 | **Ser 21** | 2.1 |
| **Phe 22** | ND | **Phe 22** | ND |
| **Glu 23** | 0.2 | **Glu 23** | 1.0 |
| **Leu 24** | 0.4 | **Leu 24** | 1.2 |
| **Phe 25** | 1.5 | **Phe 25** | 0.9 |
| **Ala 26** | 1.7 | **Ala 26** | ND |
| **Asp 27** | ND | **Asp 27** | 1.3 |
| **Lys 28** | ND | **Lys 28** | ND |
| **Val 29** | 2.0 | **Val 29** | ND |
| **Lys 31** | 2.3 | **Lys 31** | 1.2 |
| **Thr 32** | 2.1 | **Thr 32** | ND |
| **Ala 33** | ND | **Ala 33** | 1.3 |
| **Glu 34** | ND | **Glu 34** | ND |
| **Asn 35** | ND | **Asn 35** | 1.0 |
| **Phe 36** | 0.9 | **Phe 36** | 2.5 |
| **Arg 37** | 1.4 | **Arg 37** | ND |
| **Ala 38** | 0.1 | **Ala 38** | 0.4 |
| **Leu 39** | ND | **Leu 39** | 1.1 |
| **Ser 40** | 1.7 | **Ser 40** | ND |
| **Thr 41** | 0.6 | **Thr 41** | 0.5 |
| **Gly 42** | 1.3 | **Gly 42** | 2.7 |
| **Glu 43** | 0.9 | **Glu 43** | ND |
| **Lys 44** | ND | **Lys 44** | ND |

| | | | |
|---|---|---|---|
| **Gly 45** | 0.00 | **Gly 45** | ND |
| **Phe 46** | 1.3 | **Phe 46** | 0.7 |
| **Gly 47** | 2.0 | **Gly 47** | 4.7 |
| **Tyr 48** | 2.4 | **Tyr 48** | 1.1 |
| **Lys 49** | 0.6 | **Lys 49** | 0.1 |
| **Gly 50** | ND | **Gly 50** | 0.7 |
| **Ser 51** | ND | **Ser 51** | ND |
| **Cys 52** | ND | **Cys 52** | 1.4 |
| **Phe 53** | 0.5 | **Phe 53** | 2.7 |
| **His 54** | 4.0 | **His 54** | 7.2 |
| **Arg 55** | 3.4 | **Arg 55** | 3.7 |
| **Ile 56** | 0.6 | **Ile 56** | ND |
| **Ile 57** | ND | **Ile 57** | 3.2 |
| **Gly 59** | 0.0 | **Gly 59** | ND |
| **Phe 60** | 2.1 | **Phe 60** | 0.4 |
| **Met 61** | ND | **Met 61** | ND |
| **Cys 62** | 1.3 | **Cys 62** | 2.3 |
| **Gln 63** | ND | **Gln 63** | 0.0 |
| **Gly 64** | ND | **Gly 64** | ND |
| **Gly 65** | ND | **Gly 65** | ND |
| **Asp 66** | 13.4 | **Ala 66** | 0.9 |
| **Phe 67** | ND | **Phe 67** | ND |
| **Thr 68** | 18.4 | **Thr 68** | ND |
| **Arg 69** | ND | **Arg 69** | ND |
| **His 70** | 3.2 | **His 70** | ND |
| **Asn 71** | 24.5 | **Asn 71** | ND |
| **Gly 72** | ND | **Gly 72** | 2.9 |
| **Thr 73** | 2.3 | **Thr 73** | ND |
| **Gly 74** | 21.5 | **Gly 74** | 3.0 |
| **Gly 75** | ND | **Gly 75** | ND |
| **Lys 76** | 3.6 | **Lys 76** | ND |
| **Ser 77** | 10.7 | **Ser 77** | 3.9 |
| **Ile 78** | 1.2 | **Ile 78** | ND |
| **Tyr 79** | 2.7 | **Tyr 79** | 6.3 |
| **Gly 80** | 4.0 | **Gly 80** | 5.2 |
| **Glu 81** | ND | **Glu 81** | ND |
| **Lys 82** | 6.8 | **Lys 82** | 7.7 |
| **Phe 83** | 2.1 | **Phe 83** | 6.3 |
| **Glu 84** | 1.7 | **Glu 84** | 2.7 |
| **Asp 85** | ND | **Asp 85** | 2.0 |
| **Glu 86** | ND | **Glu 86** | ND |
| **Asn 87** | 4.0 | **Asn 87** | 2.0 |
| **Phe 88** | ND | **Phe 88** | 0.6 |
| **Ile 89** | 2.2 | **Ile 89** | ND |
| **Leu 90** | ND | **Leu 90** | 1.5 |
| **Lys 91** | 0.8 | **Lys 91** | 3.6 |

| | | | |
|---|---|---|---|
| **His 92** | 2.0 | **His 92** | 1.8 |
| **Thr 93** | 0.6 | **Thr 93** | 0.6 |
| **Gly 94** | 0.5 | **Gly 94** | 0.5 |
| **Gly 96** | 2.0 | **Gly 96** | ND |
| **Ile 97** | 2.1 | **Ile 97** | 3.2 |
| **Leu 98** | 3.3 | **Leu 98** | ND |
| **Ser 99** | 4.0 | **Ser 99** | 4.8 |
| **Met 100** | 1.0 | **Met 100** | ND |
| **Ala 101** | ND | **Ala 101** | 7.7 |
| **Asn 102** | 5.8 | **Asn 102** | 0.7 |
| **Ala 103** | 3.9 | **Ala 103** | ND |
| **Gly 104** | 2.2 | **Gly 104** | 0.4 |
| **Asn 106** | ND | **Asn 106** | 0.1 |
| **Thr 107** | ND | **Thr 107** | 1.3 |
| **Asn 108** | ND | **Asn 108** | 2.6 |
| **Gly 109** | ND | **Gly 109** | 1.2 |
| **Ser 110** | 5.5 | **Ser 110** | ND |
| **Gln 111** | 6.8 | **Gln 111** | 0.3 |
| **Phe 112** | 0.2 | **Phe 112** | ND |
| **Phe 113** | 4.6 | **Phe 113** | 2.6 |
| **Ile 114** | 0.8 | **Ile 114** | 0.3 |
| **Cys 115** | 0.3 | **Cys 115** | 0.5 |
| **Thr 116** | ND | **Thr 116** | ND |
| **Ala 117** | 3.1 | **Ala 117** | ND |
| **Lys 118** | 0.8 | **Lys 118** | ND |
| **Thr 119** | ND | **Thr 119** | ND |
| **Glu 120** | 1.6 | **Glu 120** | 4.4 |
| **Trp 121** | ND | **Trp 121** | ND |
| **Leu 122** | 1.8 | **Leu 122** | 1.5 |
| **Asp 123** | ND | **Asp 123** | 3.5 |
| **Gly 124** | 4.3 | **Gly 124** | ND |
| **Lys 125** | 1.8 | **Lys 125** | 0.5 |
| **His 126** | ND | **His 126** | 1.2 |
| **Val 127** | 0.6 | **Val 127** | 2.2 |
| **Val 128** | 0.8 | **Val 128** | 1.6 |
| **Phe 129** | ND | **Phe 129** | ND |
| **Gly 130** | 0.7 | **Gly 130** | 0.8 |
| **Lys 131** | 3.6 | **Lys 131** | 0.3 |
| **Val 132** | ND | **Val 132** | 2.2 |
| **Lys 133** | ND | **Lys 133** | ND |
| **Glu 134** | ND | **Glu 134** | ND |
| **Gly 135** | 1.3 | **Gly 135** | ND |
| **Met 136** | ND | **Met 136** | ND |
| **Asn 137** | 0.1 | **Asn 137** | 0.6 |
| **Ile 138** | 1.9 | **Ile 138** | ND |
| **Val 139** | ND | **Val 139** | 0.1 |

| | | | |
|---|---|---|---|
| **Glu 140** | ND | **Glu 140** | 1.7 |
| **Ala 141** | ND | **Ala 141** | 0.6 |
| **Met 142** | ND | **Met 142** | ND |
| **Glu 143** | 1.4 | **Glu 143** | ND |
| **Arg 144** | 0.1 | **Arg 144** | ND |
| **Phe 145** | ND | **Phe 145** | 0.1 |
| **Gly 146** | 1.6 | **Gly 146** | 1.9 |
| **Ser 147** | 0.8 | **Ser 147** | 1.3 |
| **Arg 148** | 1.0 | **Arg 148** | ND |
| **Asn 149** | 1.7 | **Asn 149** | 3.3 |
| **Gly 150** | 1.4 | **Gly 150** | 0.6 |
| **Lys 151** | ND | **Lys 151** | 0.0 |
| **Thr 152** | 0.0 | **Thr 152** | 1.6 |
| **Ser 153** | ND | **Ser 153** | 3.0 |
| **Lys 154** | ND | **Lys 154** | 0.3 |
| **Lys 155** | 1.4 | **Lys 155** | ND |
| **Ile 156** | ND | **Ile 156** | 2.3 |
| **Thr 157** | 0.2 | **Thr 157** | 1.1 |
| **Ile 158** | 4.1 | **Ile 158** | ND |
| **Ala 159** | ND | **Ala 159** | ND |
| **Asp 160** | 1.7 | **Asp 160** | 2.7 |
| **Cys 161** | 0.0 | **Cys 161** | 0.4 |
| **Gly 162** | ND | **Gly 162** | 1.3 |
| **Gln 163** | ND | **Gln 163** | 1.5 |
| **Leu 164** | ND | **Leu 164** | 0.5 |
| **Glu 165** | ND | **Glu 165** | 0.6 |

**Available datasets**

**Dataset S1.** 10000 representative structures of the MSM ensembles, states assignments and rate matrices for CypA, D66A, H70A CypA.

**Dataset S2:** Video summarizing the conformational changes in CypA

These datasets are available at https://bitbucket.org/jjuarez84/cypa-datasets/src

*username*:

*password*: CypA_MSM_NMR_X-Ray .
